# Desmin Mutations Disrupt Filament Elongation and Drive Polymorphic Aggregation

**DOI:** 10.64898/2026.07.22.740046

**Authors:** Lovis Schween, Jan-Philipp Burchert, Matthijs van der Heyden, Dorothea Schultheis, Norbert Mücke, Sarah Köster, Sergei V. Strelkov, Harald Herrmann, Ben Fabry

**Affiliations:** Department of Physics, Friedrich-Alexander-University Erlangen-Nürnberg, Erlangen, Germany; Erlangen Graduate School in Advanced Optical Technologies (SAOT), Erlangen, Germany; Institute for X-Ray Physics, University of Göttingen, Göttingen, Germany; Department of Pharmaceutical and Pharmacological Sciences, KU Leuven, Leuven, Belgium; Institute of Neuropathology, University Hospital of Erlangen, Erlangen, Germany; Chromatin Networks, German Cancer Research Center (DKFZ), Heidelberg, Germany; Cluster of Excellence “Multiscale Bioimaging: from Molecular Machines to Networks of Excitable Cells (MBExC)”, University of Göttingen, Germany

## Abstract

In muscle cells, desmin intermediate filaments form a cytoskeletal network that maintains the structural integrity and mechanical coupling of myofibrils. Dominant missense mutations in desmin cause myopathies characterized by intracellular protein aggregation. To explore the process of aggregate formation, we investigate the assembly kinetics of wild-type desmin and four disease-associated variants (N342D, L345P, R350P, and R406W) using dual-wavelength stopped-flow spectroscopy, complemented by atomic force microscopy and molecular dynamics simulations. At low ionic strength, all proteins form uniform tetramers. Increasing the ionic strength initiates assembly by rapid lateral association of tetramers into unit-length filaments (ULFs), followed by longitudinal elongation and radial filament compaction. R406W forms ULFs with wild-type-like kinetics, whereas N342D, L345P, and R350P exhibit delayed lateral assembly. After short filaments have formed, all four mutants diverge from productive filament maturation, but through distinct pathways. Quantitative analysis of atomic force microscopy images together with kinetic modeling of the spectroscopic data shows that wild-type filaments elongate continuously, whereas R406W filaments progressively associate into fibrillar clusters and cease elongating. By contrast, N342D, L345P, and R350P rapidly collapse into globular complexes that subsequently coalesce into larger aggregates. Molecular dynamics simulations indicate that the mutations differentially destabilize coil 2, which leads to local structural perturbations and mutation-specific assembly defects. Together, these findings identify early filament maturation - when elongating ULF-derived filaments would normally undergo radial compaction to form stable, mature filaments - as a critical time point in desmin assembly. At this stage, pathogenic mutations redirect the internal reorganization of the filament from productive stabilization toward mutation-specific structural collapse and aggregation.

**Significance:** Dominant mutations in the intermediate filament protein desmin cause myofibrillar myopathies characterized by protein aggregation and progressive muscle degeneration. Using *in vitro* filament-assembly experiments, we identify intrinsic assembly defects of disease-associated desmin mutants. Mutant proteins initially enter the normal assembly pathway, albeit with delayed kinetics. They subsequently fail at an early maturation step in which short filaments would normally reorganize to sustain productive elongation. Instead, mutant proteins form morphologically distinct aggregates. Similar structural abnormalities occur when mutant proteins co-assemble with wild-type desmin. Our findings identify the radial compaction phase during early longitudinal assembly as a critical checkpoint that determines whether desmin assembly yields functional intermediate filaments or is redirected into pathogenic aggregation pathways.

## INTRODUCTION

The mechanical phenotype of a metazoan cell is established and maintained through the dynamic organization of its cytoskeleton - an interpenetrating network of F-actin, microtubules, and intermediate filaments (IFs). The building blocks for F-actin and microtubules are actin and tubulin, which both are globular proteins. By contrast, IFs are composed of fibrous proteins (1). The defining structural feature of IF proteins is an extended central α-helical segment that mediates the formation of coiled-coil dimers (see Figure 1 A). IF proteins are highly extensible and protect cells from damage under large mechanical loads (2–7). Throughout evolution, individual members of the IF multigene family have undergone highly specific sequence adaptations to fulfill specialized functions in different tissues. This functional specialization imposes strong constraints on critical sequence elements, such that even a single amino acid substitution can severely impair IF function and cause disease, as documented for IF proteins in a wide range of tissues (8).

In muscle, numerous mutations in the muscle-specific IF protein desmin give rise to diverse forms of myofibrillar myopathy (9–11). An immunohistological hallmark of desminopathies is the presence of massive granulo-filamentous desmin-containing protein aggregates in muscle tissue. To date, more than 80 disease-causing desmin mutations have been identified (12). Notably, many of the most deleterious mutations are located within the coiled-coil forming central domain of desmin, see Figure 1 A and (13, 14). However, the impact of the individual mutations on the ability of desmin to form filaments *in vitro* exhibits a large spectrum ranging from totally blocking assembly to apparently not affecting assembly (15).

IF proteins have a tripartite structure: the central α-helical segment of ∼300 amino acids is flanked by non-structured amino- and carboxy-terminal segments of highly variant amino acid numbers (Figure 1 A). The α-helical segment (“rod”) is formed by three α-helical subsegments of 43 (coil 1A), 102 (coil 1B), and 140 (coil 2) amino acids. These three α-helical segments are connected by non-α-helical linkers L1 and L12 of slightly varying lengths between different IF proteins. Coil 2 exhibits a strict organization of periodicities: starting with three hendecad repeats, followed by six heptads, one hendecad repeat, and finally seven heptads (16). As a result of parallel coiled-coil formation of two monomers, the hendecad repeats form parallel helices, whereas the heptad repeats form left-handed coiled coils. This substructure is conserved in all classes of IF proteins, indicating it is essential for filament assembly. Therefore, mutations in the center of coil 2 and at the highly conserved IF consensus motif at the end of coil 2 are expected to be critical for coiled-coil structure and assembly.

In vitro IF assembly begins with the formation of thin, rod-like tetrameric complexes, approximately 2-3 nm in diameter (Figure 1 A, B). Each tetramer consists of two antiparallel, approximately half-staggered coiled-coil dimers. Although the individual coiled coils are ∼46 nm long, their staggered arrangement gives the tetramer a mean length of ∼60 nm (17, 18). Tetramers are regarded as the functional subunits of IF assembly (19, 20), and the conserved tetrameric organization of cytoplasmic IF proteins suggests that their subsequent assembly into elongated filaments follows a common general mechanism (21). In vitro, tetramers form spontaneously when urea-solubilized IF proteins isolated from tissues are dialyzed into low-ionic-strength buffers. Alternatively, tetramers can be generated by transferring mature IFs reconstituted from purified proteins into buffers of low ionic strength, thereby inducing filament disassembly (19).

To initiate assembly of IF protein tetramers into filaments, the salt concentration is instantaneously raised by mixing the low ionic strength tetramer buffer with a buffer of defined salt concentration in a microfluidic device (22). With this “kick-start” procedure, micrometer-long IFs form within a few minutes (23–26). Assembly proceeds in three distinct phases: (1) lateral association of tetramers into unit-length filaments (ULFs), which typically contain 8-12 tetramers (Figure 1 B); (2) longitudinal annealing of ULFs, followed by elongation through the incorporation of additional ULFs and short IFs; and (3) dynamic intra-filament reorganization during the first 10 min of assembly. This phase includes the dissociation of excess subunits from individual ULFs and radial filament compaction (27).

Together, these processes stabilize the filament and generate protofibrillar substrands, each containing two tetramers in cross-section. Protofibrils have been identified by electron microscopy (EM) as stable IF substructures, because incubation of keratin IF in phosphate buffer caused them to unravel into four distinct substrands (28). Complementary STEM mass-per-unit-length measurements indicated that keratin filaments contain 32 monomers per cross-section, corresponding to eight monomers in each substrand, or protofibril (29, 30). Moreover, depending on the assembly conditions, lateral association can generate structural polymorphism, such that filaments contain different numbers of protofibrils. Desmin filaments, for example, contain three protofibrils when assembled in 20 mM salt but six protofibrils when assembled in 100 mM salt (31).

The first phase of the assembly process from tetramers to ULFs progresses rapidly so that most free tetramers and octamers have mostly disappeared by 50 milliseconds, consistent with a diffusion-limited lateral association process (19). The kinetics of the second longitudinal annealing phase depend more sensitively on the ionic conditions and differ among IF types. The intra-filamentous reorganization phase starts during the longitudinal assembly phase and is largely complete after 10 minutes of assembly (27, 32). At longer time scales, IF length reaches an equilibrium, as very long filaments elongate less frequently and may fracture stochastically (33).

**Figure 1.**
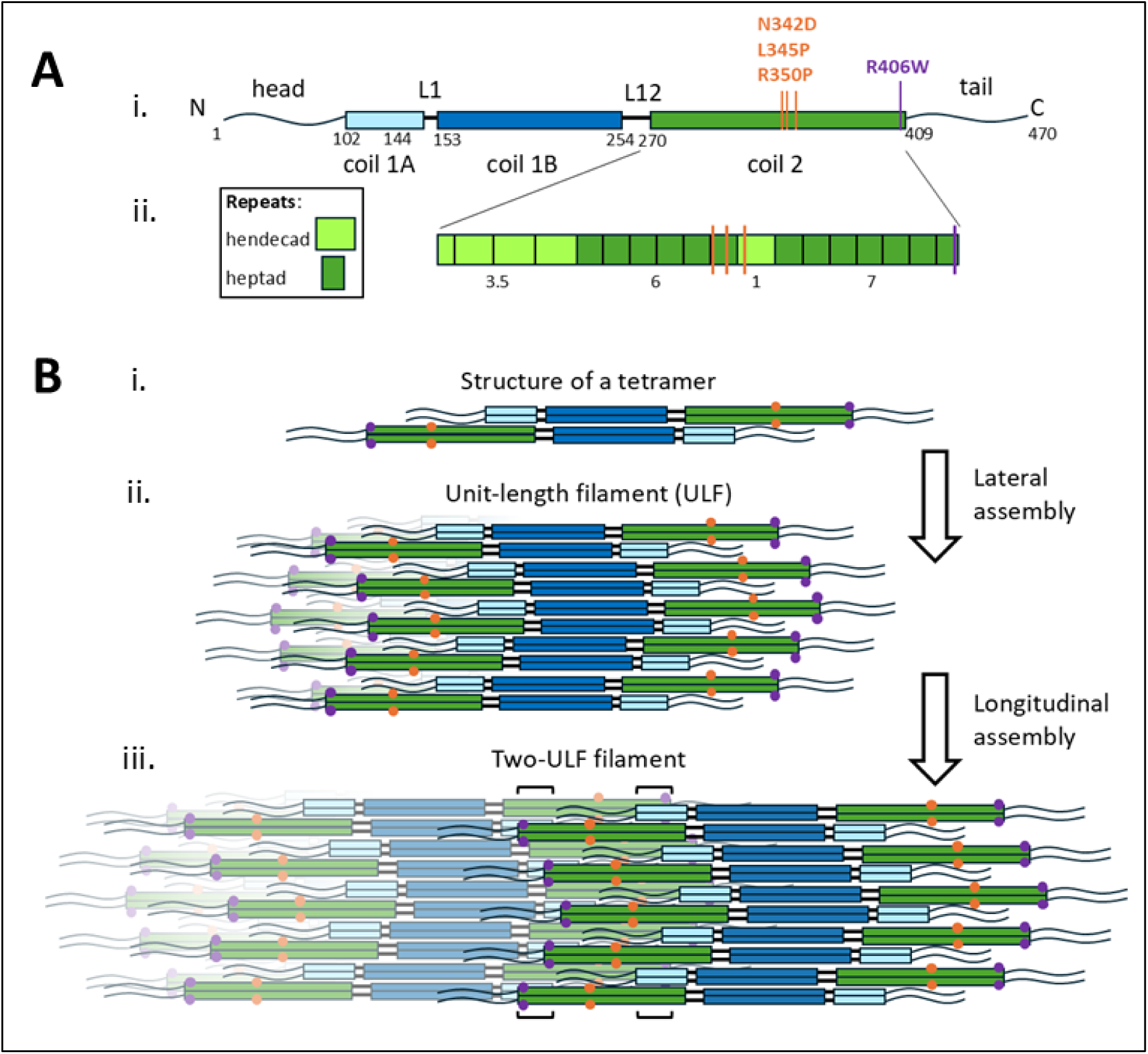
Structure and assembly mechanism of intermediate filament proteins. **(A)** Schematic overview of the desmin monomer structure. (i) Boxes represent α-helical segments connected by non-α-helical linkers L1 and L12, referred to as rod; head and tail depict the intrinsically disordered amino and carboxy terminal domains. N, amino terminus; C, carboxy terminus. Numbers indicate the amino acid positions of the respective domain borders. The positions of the mutated amino acids are indicated by dashes; they are presented in the single letter code with the wild-type amino acid before the position number, followed by the mutant amino acid. (ii) Substructure of coil 2: hendecad repeat, bright green box; heptad repeat, dark green box. The mutations N342D and L345P are in front of the single hendecad segment, R350P is within the hendecad segment. The R406W mutant is located in the last heptad repeat of coil 2. **(B)** Assembly scheme of IF proteins. (i) The tetramer is the key element for filament assembly and consists of two coiled-coil dimers arranged in an antiparallel, half-staggered manner. (ii) Phase one: lateral association of eight tetramers to a unit-length filament (ULF). (iii) Phase two: For filament elongation, two ULFs anneal via an overlap interaction of the end segments of coil 1A of the one ULF with coil 2 of the second ULF. The overlap is about 4 nm, indicated by the brackets. The mutations in the center of coil 2 (light orange dots), are in close contact with the mutations in the interacting coil 2 dimer of the neighboring ULF (dark orange dots). The R406W mutation of each ULF is in contact with coil 1a of the neighboring ULF (purple dot at the end of coil 2).

For the analysis of the IF assembly, a number of complementary methods have been employed: electron microscopy (EM), atomic force microscopy (AFM), small-angle X-ray scattering (SAXS), rheology, and hydrogen-deuterium exchange mass spectroscopy (26, 35–47). With these techniques, the assembly kinetics can be measured in a time window from typically 1 s to 30 min (23, 48). However, they cannot resolve sub-second kinetic processes due to sampling constraints, or processes over longer time scales due to filament entanglement. Light scattering techniques and total internal reflection microscopy have been used to follow the assembly from several seconds to hours (24, 26). Moreover, light scattering in combination with a stopped-flow approach can resolve kinetic processes during the very early phases of assembly, with a time resolution of 1 ms (19). By implementing dual-wavelength stopped-flow spectroscopy (DWSF), the simultaneous measurement of growing filaments with two wavelengths can distinguish between kinetic processes of the lateral versus longitudinal assembly, as the scattering behavior of light at different wavelengths sensitively depends on the shape of the assembly products (49, 50).

We examined the assembly kinetics of wild-type desmin and four desmin mutants responsible for severe disease. These mutants diverge from productive filament formation and undergo extensive aggregation in a mutant-specific manner. Although the associated phenotypes have been documented previously, this has only been done qualitatively using time-lapse electron microscopy (15, 51–53). Here, we aim to elucidate in a quantitative manner at which time point of the assembly process these mutants leave the regular assembly pathway, and at which speed they form aggregates. To analyze the scattering data, we model the assembly products as worm-like chains with different persistence lengths, or as 3D clusters with different fractal dimensions (54–57). To validate our modeling approach, we also measure the shape of the assembly products with AFM at different time points. We find that the lateral assembly of wildtype and mutant desmin proceeds with a similar kinetics, although three mutants start lateral association only after a lag phase. Moreover, the different mutants exhibit vastly different kinetic processes during the lateral assembly phase that give rise to distinct aggregate morphologies.

## MATERIALS AND METHODS

### Protein purification, reconstitution and assembly

Desmin is expressed in *E. coli*, isolated from inclusion bodies and purified by column chromatography as previously described (22, 37). The protein is solubilized in 8 M urea and is renatured by a stepwise dialysis from 8 M with steps of 2 M urea to urea-free low ionic strength buffer, referred to as tetramer buffer. For our experiments we use two tetramer buffers: 5 mM Tris-HCl (pH 8.4) and 2 mM MOPS (pH 7.5). In addition to the published procedure, we have introduced a purification step: During stepwise renaturation of the recombinant protein, we perform the 4 M urea dialysis overnight using dialysis tubing with a cut off of 50 kDa (Spectra-Por 6 pre-wetted RC tubing, Fisher Scientific, Schwerte, Germany), followed by further dialysis steps into tetramer buffer. This additional step removes minor molecular weight components including bacterial proteases. The purity and stability of the protein is controlled by sodium dodecyl sulfate polyacrylamide gel electrophoresis (Figure S1). A protease-free preparation is important for stopped-flow experiments that are performed over the course of several days after renaturation. In our preparations, degradation is completely absent as documented by the absence of additional bands on the gel after one day of dialysis (Figure S1, lanes labeled a). Because the unstructured head and tail domains are more protease-sensitive than the rod domain, proteolysis would be expected to produce a 43 kDa fragment corresponding to the rod domain (22, 37, 58). For the experiments, mouse and human desmin are used, which only deviate in 11 homologue amino acid exchanges and exhibit highly similar assembly behavior (Figure S2).

For DWSF and AFM experiments, the protein concentration is determined by Bradford protein Assay (Coomassie Plus Protein Assay; Thermo Fisher Scientific, Rockford, Il, USA). Assembly is initiated by adding an equal volume of a two-fold concentration of the respective salt to a two-fold concentration of the protein, both in tetramer buffer. The final protein concentrations are 0.1 and 0.2 mg/ml (corresponding to 0.5 and 1 µM tetramers, respectively) to minimize filament entanglement. The DWSF and AFM experiments are carried out at 37 °C.

For SAXS experiments, the protein concentration is determined with a micro-UV/Vis spectrophotometer (NanoDrop OneC, Thermo Fisher Scientific) at 280 nm, with a 340 nm baseline correction and UVettes (Eppendorf, Germany). The respective protein concentration ranges from 0.2 mg/ml to 1.25 mg/ml desmin. The majority of measurements are done with 1.2 mg/ml human, and 1.25 mg/ml mouse desmin. Assembly is initiated by adding 10X salt solution to the protein solution at a 1:9 (v/v) ratio, followed by transfer into capillaries. Samples are incubated for 4 h at 37 °C. Measurements are subsequently acquired at 10 min intervals over 12 h at room temperature.

### Dual-wavelength stopped-flow spectroscopy

The dual-wavelength stopped-flow spectroscopy (SF-61, Hi-Tech Scientific, Salisbury, UK) setup is used as described previously (19, 50). For all experiments, thoroughly degassed buffers are used. The samples are filtered through 0.22 µm sterile filters (Rotilabo PVDF filter, Carl Roth, Karlsruhe, Germany) immediately before use. Assembly conditions are identical to those previously established for vimentin (50). Scattered light intensities are sampled at 1 kHz and digitized at 12-bit resolution. Kinetic measurements are repeated three to six times consecutively and combined into a median intensity profile for each assembly condition. These scattered light intensity profiles are baseline-corrected by subtracting the protein-free buffer baseline. However, a drift in the protein-free buffer baseline during sequential measurements on the same day, despite rigorous degassing and filtration of all solutions, imposes a persistent challenge. To mitigate this effect, we use the fact that intensity ratio of the scattered light of different wavelengths, I_594_/I_405_ must be constant for the tetrameric scatterers at assembly start. This allows us to fit the drifting baseline and normalize the profiles to unity within the first 100 ms. The resulting traces are smoothed using a Butterworth low-pass filter with a cutoff of 400 Hz.

For every experiment, the proteins are freshly renatured. For each desmin variant and salt condition, 3-20 independent measurements are performed. The profiles shown in Figures 4, 6, 9, and 10 represent normalized median traces from 3-6 runs of a single representative experiment. An overview of all measured scattering intensity profiles and the corresponding intensity ratios is provided in Figure S3 for 0.1 mg/ml desmin and in Figure S4 for 0.2 mg/ml wild-type desmin.

### Atomic force microscopy

Atomic force microscopy (AFM) is performed in tapping mode in air using an Asylum Research Atomic Force Microscope MFP-3D (Asylum Research, Santa Barbara, CA, USA) and 125 µm long tips with 40-48 N/m spring constant (Tap190AL-G, BudgetSensors NanoAndMore GmbH, Wetzlar, Germany; SSS-NCHR-SPL, BudgetSensors NanoAndMore GmbH, Wetzlar, Germany) as previously described (59). For AFM, filament assembly experiments are carried out in a volume between 300 µl and 500 µl in Eppendorf tubes. The assembly process is stopped and the sample fixed by a 3-or 5-fold dilution in the same salt solution containing 0.25% glutaraldehyde (Merck, Darmstadt, Germany) for 1 min. 50 µl of the fixed filament solution is then deposited on freshly cleaved mica. After 1-3 min, the mica is washed with double-distilled water and dried with a filtered, low-pressure air stream. Imaging is done in AC Air Topography mode using Igor Pro 6.37 (WaveMetrics, Portland, OR, USA). The height and amplitude images are contrast-enhanced by percentile contrast stretching and converted to PNG format using Python. The filaments are tracked manually using ClickPoints (Version 8.0.4) (60).

### Small-angle X-ray scattering

Small-angle X-ray scattering (SAXS) is performed using an in-house setup (Xeuss 2.0, Xenocs, Sassenage, France) as described previously (40, 44). The X-radiation is produced by a Cu K_α_ source (*E* = 8 keV, Genix 3D, Xenocs) operated at 50 kV and 600 µA, with multilayer optics and scatterless slits. The beam size at the sample is approximately 0.5 x 0.5 mm^2^, with an incident intensity of approximately *I_0_* ∼8.5 x10^6^ counts s^-1^. Scattering is recorded using a Pilatus3 R 1M detector (Dectris Baden, Switzerland; 981 x 1043 pixels, pixel size 172 x 172 µm^2^) at a sample-to-detector distance of 1218 mm, providing a range of 0.08 < *q* < 5 nm^-1^. Data reduction, buffer subtraction, normalization and intensity calibration were performed as described previously (44). The cross-sectional radii of gyration *R_C_* are determined from the individually processed scattering profiles using the rod-shaped-particle Guinier analysis implemented in PRIMUS (ATSAS v3.1.1) (61):

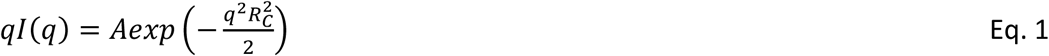

Accordingly, *R_C_* is obtained by linear regression of *ln*[*qI*(*q*)] against *q^2^*over profile-specific intermediate-*q* intervals individually for every measurement, satisfying *q*_max_*R_c_* < 1.0 (Figure 2 B, Figure S5). The selected fitting intervals are inspected for linearity (Table S1). R_C_ describes the cross-sectional rather than the overall radius of gyration of the filament. Assuming a homogeneous circular cross-section, apparent filament diameters can be estimated as 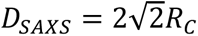 (Table S1).

### Worm-like chain (WLC) simulation

Filaments are modeled as worm-like chains (WLC) of contour length *L = N*a*, discretized into *N* identical beads with a fixed contour spacing of *a* = 12.5 nm. The scattered light intensity of a WLC is modeled by summing the scattered light intensity contributions of the individual beads and the scattered light interference terms of all possible bead pairs along the chain, following the approach of Debye (62). Each bead produces a complex scattering amplitude *A_λ_* that depends on the refractive-index contrast *Δn*, bead volume *V*, wavelength *λ*, and form factor *P*:

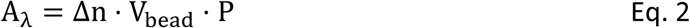

For spherical particles of radius R, the form factor is given by Rayleigh-Gans theory according to 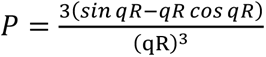, with light scattering vector 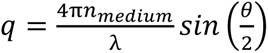 according to and scattering angle θ = 90°, following our experimental setup (49, 63). The scattering vector calculated for both wavelengths yields 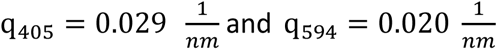. The total scattering amplitude of a WLC is given by the sum of the amplitudes of each bead i along the WLC according to

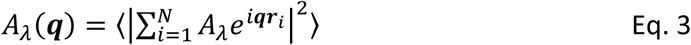

Finally, the scattered light intensity is obtained as the ensemble-averaged squared modulus of this amplitude and an interference term for all bead pairs *i, j* along the chain according to

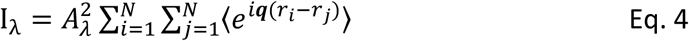

where *r_i_ (r_j_)* denotes the position of bead *i* (bead *j*) along the chain. The squared total scattering magnitude is ensemble-averaged over all WLC conformations and isotropic filament orientations.

As the WLC is statistically homogenous along its contour, the interference term 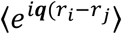 depends only on the contour separation *s = k*a*, with separation index *k*=|*j* − *i*| and bead spacing *a* = 12.5 nm. Therefore, it can be written in the form of Debye (62)

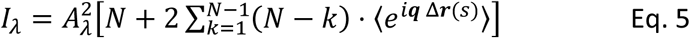

with Δ**r**(s) being the Euclidean distance between two beads separated by contour distance s. The interference term 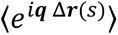 is approximated as Gaussian distribution with variance 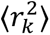 as

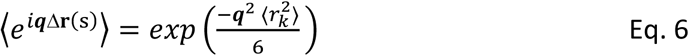

where 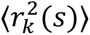 depends only on the contour length s and persistence length *L_p_*

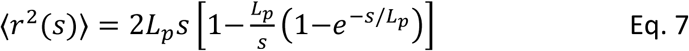

This Gaussian approximation accounts for a distribution of bead-bead separations and therefore eliminates single high-frequency interference effects. It reflects the polydispersity of WLCs as seen also in the different shapes and sizes of mutant desmin aggregates. Taken together, we can simulate the wavelength-dependent scattered light intensity of a WLC with arbitrary contour length L and persistence length *L_P_*.

### Dimensionality analysis

Unit-length filaments (ULFs) are approximated as spherical scatterers of radius a_0_ = 16.5 nm, which corresponds to the volume equivalent of a single ULF (∼10 nm x 60 nm). ULFs are considered to form self-similar clusters with prescribed fractal dimensions. The fractal dimension D_f_ is a unitless measure of the internal packing density, defined by scaling the structure mass with size. A fractal dimension of unity corresponds to a single-stranded elongated filament; a fractal dimension of 3 corresponds to a densely packed sphere. The number N_ULF_ of ULFs contained within a fractal cluster of radius *R* ≥ *a_0_* scales with R and D_f_ according to (54)

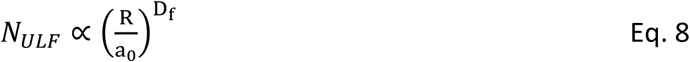

The scattered light intensity of a single cluster increases with the product of the number of ULF-equivalent scatterers *N_ULF_*, the single-ULF form factor *P_ULF_(q),* and a size-dependent fractal structure factor *S(q, R):*

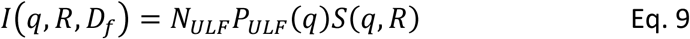

The single-ULF form factor *P_ULF_(q)* describes the scatter intensity of isolated ULFs and is approximated by a Guinier-type expression:

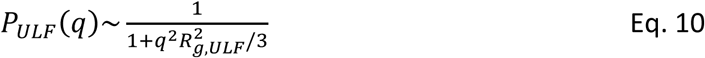

where R_g,ULF_ is the radius of gyration of a single ULF. The wavelength-dependent scatteringvectors q are 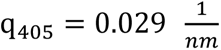 and 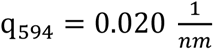, as derived above. The fractal structure factor *S(q, R)*, also called Teixeira-factor, depends on *q, R* and D_f_ according to (55)

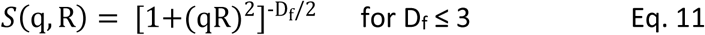

For small cluster (*qR* ≪ 1), corresponding to *R* << 33 nm at 405 nm and R << 50 nm at 594 nm, the fractal structure factor remains unity and the scattering is dominated by the single ULF form factor *Pulf*. In this regime, the scatter intensity is effectively cluster size-independent, and the simulated intensity ratios I_594_/I_405_ remain constant, consistent with Rayleigh scattering (49).

As clusters grow to the sizes comparable to or larger than the incident wavelengths (*qR* ≫ 1), the wavelength-dependence introduced by the structure factor (Eq. 11) causes the intensity ratio to converge toward a fractal-dimension-dependent plateau (54, 55):

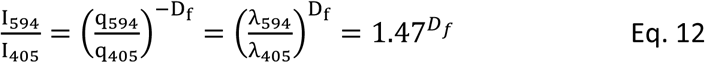

### Simulation of morphology

Desmin filaments are modeled as worm-like chains of appropriate persistence lengths. Contour lengths are drawn from a log-normal distribution reflecting the experimentally observed polydispersity. To recapitulate the shape of the WLC aggregates as observed by AFM imaging, the WLCs contours are projected onto the xy-plane and convolved with a 2D Gaussian kernel approximating the AFM tip radius, with hillshade-based contrast rendering to replicate AFM appearance. Protein aggregates are simulated using a fractal growth algorithm with fractal dimension D_f_ as the single morphological control parameter. All simulations are performed using custom Python scripts.

### Molecular dynamic simulations

The starting wild-type desmin coil 2 model is based on a previously reported crystal structure (PDB code 1gk4) of the corresponding vimentin fragment (18). The pathogenic mutations are modelled in COOT (64). All MD simulations are performed using Desmond (Schrödinger suite 25-2 release) (65). Protein systems are prepared in Maestro (Schrödinger) using the Protein Preparation Wizard, including assignment of protonation states and capping of terminal residues (66). Systems are solvated using the TIP4P water model in an orthorhombic simulation box with a 15 Å buffer in each dimension and supplemented with 0.15 M NaCl to mimic physiological conditions. The final system sizes range from approximately 100,000 to 200,000 atoms.

To explore the preservation of the α-helical structure, the simulation is carried out for 100 ns at 300 K using the OPLS4 force field, with trajectory frames recorded every 25 ps (67). To maintain structural stability of the N-terminal region, residues 268–307 are restrained with a force constant of 25 kcal/(mol¹·Å²). To explore the ability of the MD to recover the correct coiled-coil conformation, the model is manually unzipped at residue 328 using COOT prior to the MD simulation.

## RESULTS

To perform kinetics measurements with dual-wavelength stopped-flow spectroscopy (DWSF), we use a “kick-start” mode for filament assembly by raising the salt concentration (13, 15, 31, 68). Since salt concentration and buffer type influence IF assembly in a protein-specific way (69), we first evaluate the impact of salt concentration on desmin filament assembly and morphology.

### Small-angle X-ray scattering of desmin filaments

Small-angle X-ray scattering (SAXS) measures the morphology of unfixed filaments in solution. From the scattered radial intensity as a function of the scattering vector (Figure 2 A), we compute the cross-sectional radius of gyration of the assembled filaments for mouse and human desmin (Eq. 1; Figure 2 B). Without salt, no filaments but only tetrameric complexes are formed, as previously shown by analytical ultracentrifugation (15, 38). In this case, the resulting cross-sectional radius of gyration (R_C_) of mouse desmin is with 2.90 ± 0.13 nm lower than the R_C_ of human desmin with 3.49 ± 0.2 (Figure 2 B, Table S1). With increasing salt concentration in 20 mM steps up to 100 mM, the scattering signals at small scattering vectors increase monotonically. Beyond 40 mM, this increase becomes more gradual (Figure 2 A). The resulting cross-sectional radius of gyration becomes more similar for mouse and human desmin and are between 4.9 nm at 40 mM and 5.6 nm at 100 mM (Figure 2 B; Table S1). The corresponding filament diameters D are between 13.7 nm at 40 mM and 15.8 nm at 100mM. The similarity of data for mouse and human desmin filaments reflects the high identity of both amino acid sequences. Moreover, the diameters obtained are in reasonable agreement with previous scanning transmission electron microscopy (STEM) measurements, which reported an increase from 9.5 nm at 20 mM salt to 12.6 nm at 100 mM (31), and values obtained by conventional EM with 13.9 and 13.5 (27, 38).

STEM and EM principally yield lower values due to the fixation and deposition of filaments onto a solid support, followed by air-drying. Furthermore, in negatively stained TEM images, the filaments outlines are manually selected, from which the diameter is measured. STEM avoids this ambiguity by measuring the full width at half maximum of the transverse mass profile, which necessarily yields a smaller value than the outer diameter. When instead the filaments outlines were manually selected from STEM images, diameters of 15-16 nm were obtained (70), which is in agreement with our SAXS results.

**Figure 2.**
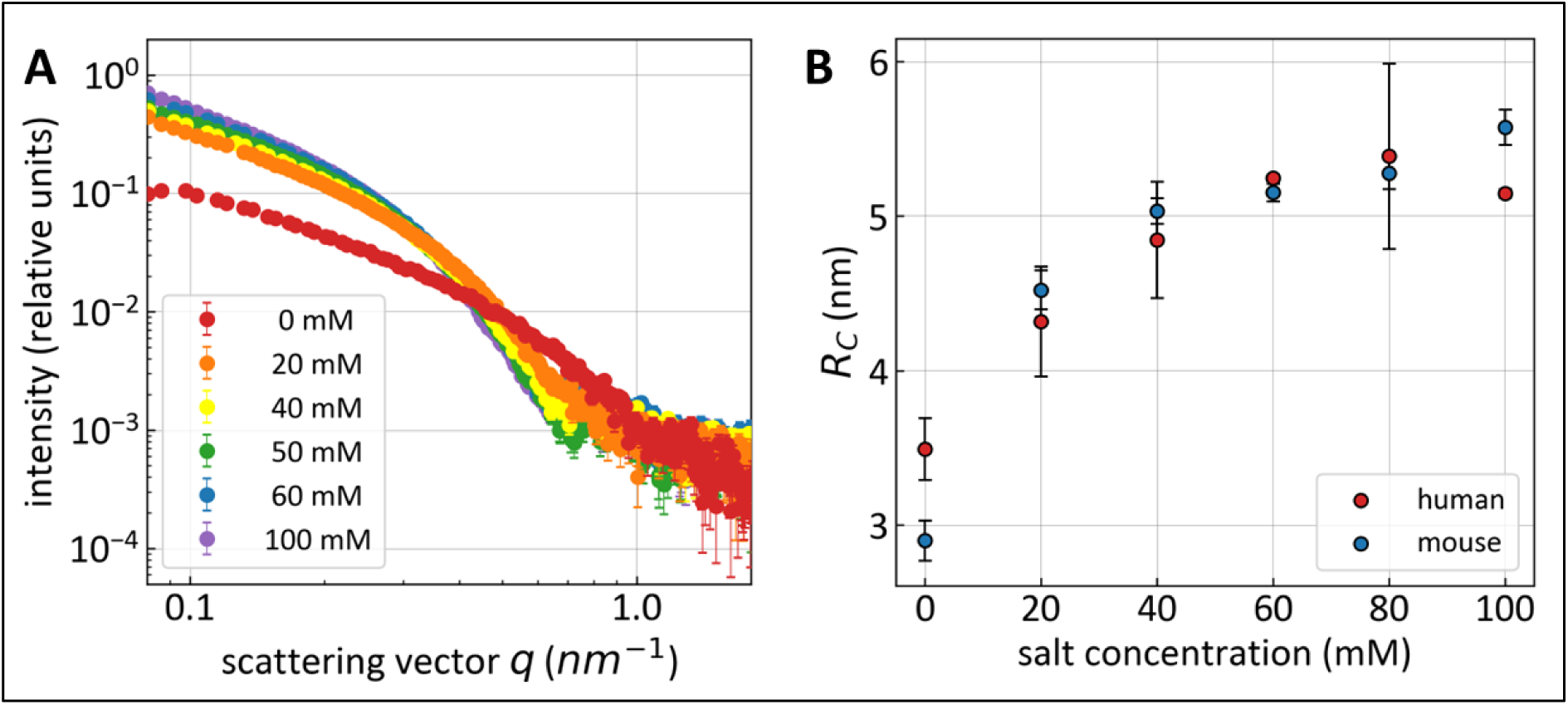
Small-angle X-ray scattering of desmin filaments. **(A)** Scattering profiles wild-type mouse desmin (1.25 mg/ml) assembled at the indicated KCl concentrations in 2 mM MOPS, pH 7.5. Intensities were normalized to the protein concentration and are proportional to the differential scattering cross-section per unit mass. Symbols and error bars represent mean ± SD; at low *q*, the error bars are smaller than the symbols. **(B)** Cross-sectional radii of gyration RC obtained by cross-sectional Guinier analysis for mouse desmin (blue) and human desmin (red) assembled under the same buffer and salt conditions. Symbols and error bars represent mean ± SD from two to nine individual measurements per salt concentration and species. Data provided in Table S1, Guinier fits displayed in Figure S5.

### *In vitro* assembly of wild-type desmin filaments

First, we characterize the filament formation process by measuring the mean length and length distribution of the desmin filaments after 1 min, 5 min, and 10 min of assembly at 50 mM salt (Figure 3 A-C). The measured filament length is expressed as the number of ULF-segments per filament. One ULF corresponds to a length of 60 nm (35). Each additional ULF adds 43 nm due to the overlap between adjacent ULFs (25, 48). We find that the desmin filament lengths are log-normal distributed (Figure 3 D-F). A log-normal distribution has previously been reported for vimentin filaments as well (50).

**Figure 3.**
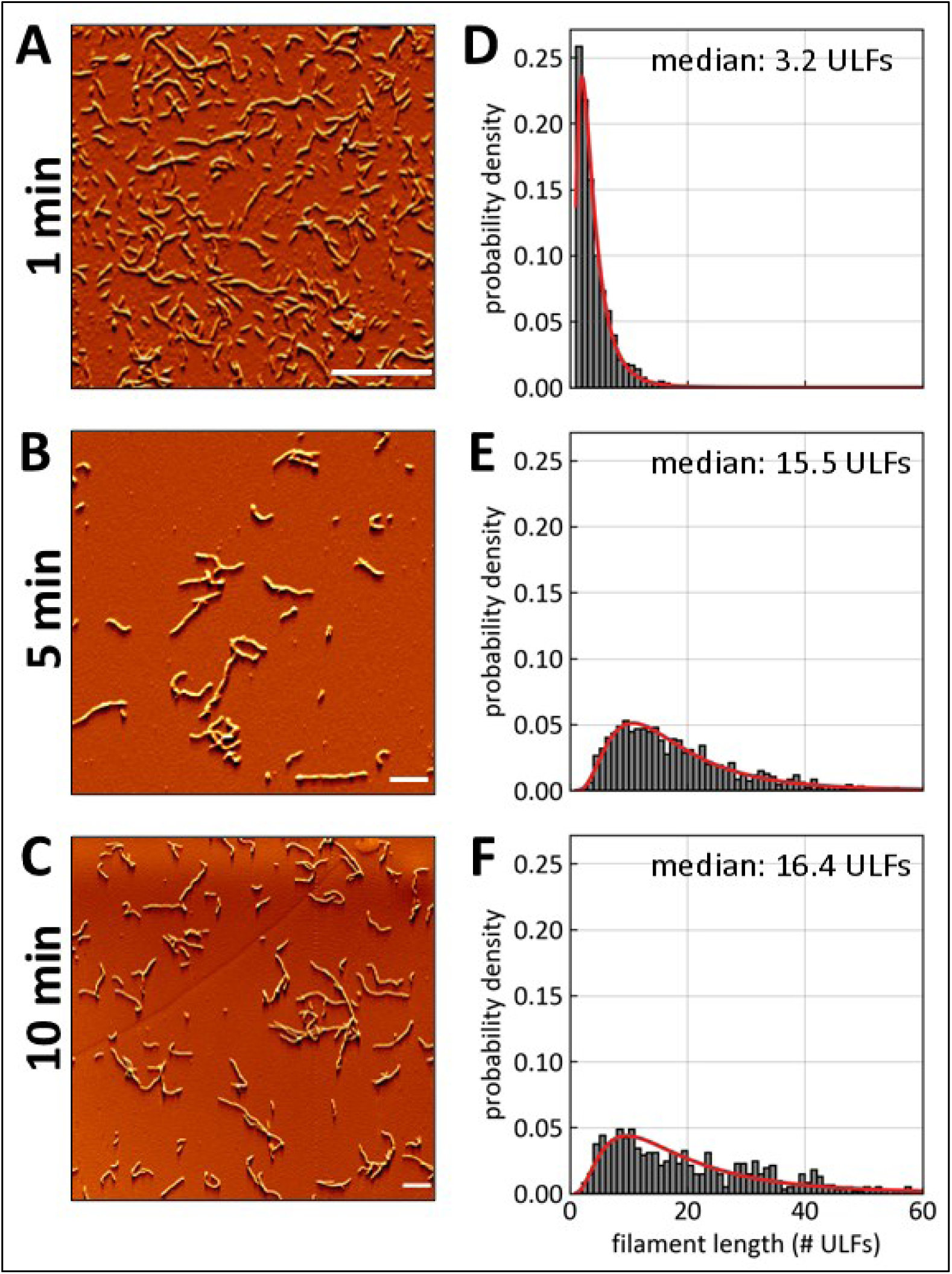
Atomic force microscopy of desmin filaments. **(A-C)** Atomic force microscopy images of 0.2 mg/ml wild-type desmin assembled for 1 min, 5 min, and 10 min in 2 mM Mops buffer, pH 7.5, with 50 mM KCl at 37°C. Bar: 1 μm. **(D-F):** Probability density distribution of desmin filament lengths (in units of #ULF). Bin width corresponds to 1 ULF. Between 620 to 1770 filaments are analyzed per condition. The red line shows the fitted log-normal distribution; the median filament length corresponds to the geometric mean of the distribution and is specified in each figure in units of #ULFs.

The average desmin filament length increases from 3 ULFs ± 0.7 (geometric mean ± standard deviation) after 1 min to 15 ULFs ± 0.6 after 5 min. This increase corresponds to an average elongation rate of approximately 3 ULFs per minute, which is a 3-fold higher than that of vimentin (50). At 10 min of assembly, the median filament length growth appears to slow down (16 ULFs ± 0.7). However, this is largely attributable to preferential entanglement of longer filaments, which prevents their lengths from being reliably determined in AFM images.

### Dual-wavelength stopped-flow spectroscopy of desmin assembly

To study the kinetics of filament assembly with high time resolution, we use DWSF spectroscopy, as previously established for vimentin (50). Static light scattering is performed at an angle of 90°. The filament assembly is initiated in a reaction chamber of the stopped-flow apparatus by mixing equal volumes of desmin tetramer solution and salt solution within 3 ms. The final concentrations are 50 mM or 100 mM salt and 0.1 mg/ml desmin. The assembly process is followed by recording the scattered light intensities at two wavelengths (405 nm and 594 nm) over time. The scattered light intensities for each wavelength are normalized to the corresponding scattered light intensity of the tetramer solution. The intensities report the average molecular weight of the protein complexes relative to tetramers as long as their size remains within the Rayleigh scattering regime, which is fulfilled by the single ULF (50).

When we compare the desmin assembly data with previously published data for vimentin (50), another type III intermediate filament that shares the same three-phase assembly pathway, we find a similar characteristic pattern: an initial synchronous rise of both wavelength signals, a shoulder that precedes the longitudinal assembly, and a subsequent divergence of the 405 nm and 594 nm signals as filaments elongate (Figure 4 A, B). The shoulder marks a stage at which rapid ULF formation is nearly complete, and the onset of ULF elongation has not yet produced a detectable increase in scattered light intensity. Higher salt concentrations accelerate both, the lateral and even stronger the longitudinal filament assembly.

In the following, we use the occurrence of the shoulder, which marks the completion of the lateral assembly phase, to quantify the lateral assembly kinetics. At 50 mM salt, the shoulder occurs at 730 ± 130 ms (mean ± standard error). At 100 mM salt, the scattering signals show a qualitatively similar pattern, but with steeper slopes and a less pronounced shoulder that occurs at 430 ± 70 ms (Figure 4 A, B, Table S2), indicating a ∼45% shorter lateral assembly phase compared to 50 mM. The higher signal intensity of the shoulder measured for the 100 mM conditions compared to 50 mM can be explained by a higher number of tetramers per ULF, since the diameter of the filaments, and therefore the mass-per-cross section, increases with the salt concentration used for assembly (Figure 2). Furthermore, the less pronounced shoulder at 100 mM compared to 50 mM salt is explained by a reduced temporal separation between the lateral and longitudinal assembly phases.

Compared to desmin, vimentin assembly at 50 mM salt exhibits a slower increase in intensity during the lateral assembly phase (Figure 4 A, B) but a similar timing of the shoulder (760 ± 350 ms), indicating that the lateral assembly kinetics of desmin and vimentin are comparable. At 100 mM salt, however, lateral assembly completes nearly 40 % faster in vimentin (250 ± 50 ms) than in desmin (430 ± 70 ms). The slower lateral assembly kinetics in desmin may arise from its larger number of tetramers per ULF (12 tetramers at 100 mM) compared to vimentin (8 tetramers) (27, 31). Additionally, the lateral assembly of desmin is less salt dependent than that of vimentin.

When filaments elongate, wavelength-dependent form factor effects suppress the rise of the 405 nm signal more strongly than the 594 nm signal, as described by Rayleigh-Gans theory (Figure S6). Therefore, the two signals diverge when a substantial fraction of the filaments reaches a length comparable to 2 ULFs or more (50). Consequently, the intensity ratio I_594_/I_405_ increases with increasing filament length and allows us to dissect the kinetics of the lateral and longitudinal assembly (50).

For desmin, the intensity ratio I_594_/I_405_ begins to rise shortly after the occurrence of the shoulder in the scattering curves, whereas for vimentin, the rise of the intensity ratio is delayed (50mM: 19 ± 4 s in vimentin vs 1.4 ± 0.1 s in desmin, 100 mM: 2.2 ± 0.4 s in vimentin vs. 1.0 ± 0.03 s in desmin; Figure 4 C, D) . This faster rise in desmin indicates a faster longitudinal assembly kinetics compared to vimentin.

The intensity ratios follow, on a linear time axis, a saturating approximately exponential profile (Figure S7). We fit a simple exponential function *f*(*x*) = 1 + *A*(1 − *e*^−*t*/*τ*^) to the data and extract the time constant τ as a quantitative measure of the elongation process. For desmin, the mean ± SE time constants are 76 ± 11 s at 50 mM salt and 57 ± 7 s at 100 mM, indicating a slightly faster elongation at higher ionic strength. For vimentin, the respective time constants are 258 ± 52 s and 120 ± 18 s, revealing a stronger salt-dependence but overall slower kinetics (50). The time constants τ, expressed in minutes, are converted into filament growth rates using the relation *r_elong_* = 2.32 / τ in ULFs/minutes (50). Based on light scattering data obtained for 0.1 mg/ml desmin at 50 mM salt, desmin filaments elongate at a rate of 1.8 ULFs/min. This is approximately 3.5 times faster than the rate of 0.5 ULFs/min observed for vimentin. At 100 mM salt, desmin filaments elongate at a rate of 2.4 ULFs/min, which is approximately twice as fast as vimentin. When the same conversion of τ in r_elong_ is applied to the protein and salt concentrations used for AFM imaging (0.2 mg/ml desmin and 50 mM salt, Figure 3), the resulting DWSF-derived growth rate is 3.2 ULFs/min (Figure S7), which closely matches the growth rate derived by AFM.

**Figure 4.**
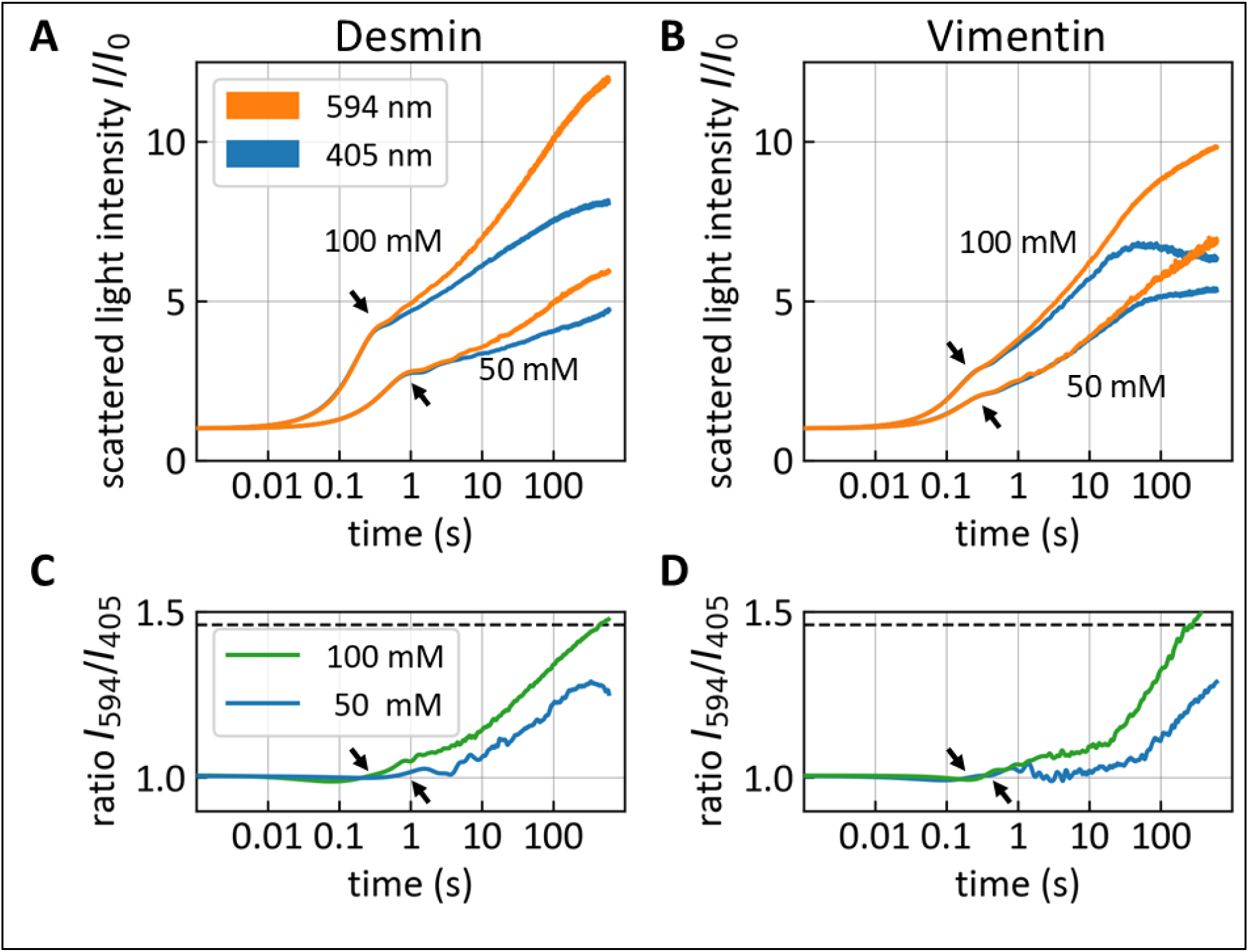
Assembly kinetics of desmin and vimentin measured with DWSF spectroscopy. **(A, B)** Scattered light intensity profiles of desmin (A) and vimentin (B) as measured by DWSF light scattering. Intensities are recorded at 594 nm (orange) and 405 nm (blue). Assembly of 0.1 mg/ml desmin (A) and vimentin (B) assembled at 50 mM and 100 mM sodium chloride as indicated in the figure. The median of five consecutively taken measurements per condition is shown. **(C, D)** Intensity ratios I594 /I405 of desmin (C) and vimentin (D) of data shown (A) and (B), respectively. Arrows: Inflection point in the scattering light intensities profiles of the two wavelengths. Vimentin data (B, D) are from (50).

### Molecular-dynamics simulation of desmin and mutated desmins

The wild-type amino acids at the mutated positions analyzed in this study (N342, L345, R350 and R406) are highly conserved across all IF proteins (Figure S8). Within the IF sequence homology class III (vimentin, desmin, GFAP and peripherin), all these residues are identical. In neuronal IF proteins, sequence homology class IV, leucine 345 is fully conserved, while positions 342 and 350 show conservative substitutions (Figure S8 A). Comparisons of desmin sequences across vertebrates—from sharks to humans—reveal strong conservation at these positions, where even distant chordates like the sea lamprey and pleated sea squirt retain remarkable similarity (Figure S8 B). Although regions located downstream of R350 vary more with evolutionary distance, the final ∼30 amino acids, including the IF consensus motif, are almost completely conserved. This high degree of conservation in this region appears to be linked with their involvement in the head-to-tail dimer-dimer interactions during desmin filament elongation. In these interactions, the coil 1A segments of one dimer form an approximately 4-nm parallel overlap with the C-terminal coil 2 segments of another dimer, thereby maintaining a uniform N-to-C orientation along each longitudinal dimer chain (Figure 1 B iii).

To investigate the importance of the mutated residues for the structural properties and the dynamics of coil 2, we perform *in silico* modeling using AlphaFold3. No significant differences are observed in the predicted structures for any of the variants, all of which are predicted to form an α-helical, parallel coiled-coil dimer (Figure 5 A and Figure S9).

We then run molecular dynamics (MD) simulations on the wild-type sequence of coil 2 (residues 268–414) and its respective mutants. Specifically, we analyze whether the α-helical structure was preserved during simulation (Figure 5 B). In the wild-type protein, coil 2 remains predominantly α-helical, with only a minor destabilization at the stutter region, which is the single hendecad repeat indicated in Figure 1 A. Additionally, the final five residues lose their α-helicity, consistent with the coil 2 terminating at residue 409 (16, 71).

Analysis of the mutants reveals distinct effects. The L345P and R350P mutations cause a clear, localized loss of helicity at the mutation sites, consistent with the helix-disrupting nature of proline residues (72). These disruptions may facilitate the formation of flexible ‘linker-like’ regions. By contrast, the N342D and R406W mutations do not induce detectible changes in helicity in these simulations (Figure 5 B).

Next, we use MD to explore whether the wild-type and respective mutant structures return to the correct coiled-coil conformation after artificial unzipping. To this end, coil 2 is manually unzipped from residue 328 onwards for all structures (except for R406W as it effectively locates at the very C-terminus of the structure) (Figure 5 C). The resulting partially splayed helices are then subjected to MD simulations. Here, the wild-type coil 2 rapidly (within 5 ns) recovers the native coiled-coil dimer structure (Figure 5 C). By contrast, the N342D mutant shows incomplete reassembly, with helix association stalling at the mutation site and preventing restoration of the native coiled-coil structure. The L345P variant can reassociate as well; however, loop-like distortions form near the introduced proline residue and are more pronounced than in the preceding simulations. Similarly, the R350P mutant arrests at the mutation site, where proline-induced kinking prevents close packing of the helices. Although the helices approach each other within 60–80 ns in all mutant simulations, correct coiled-coil formation is not recovered for N342D or R350P.

**Figure 5.**
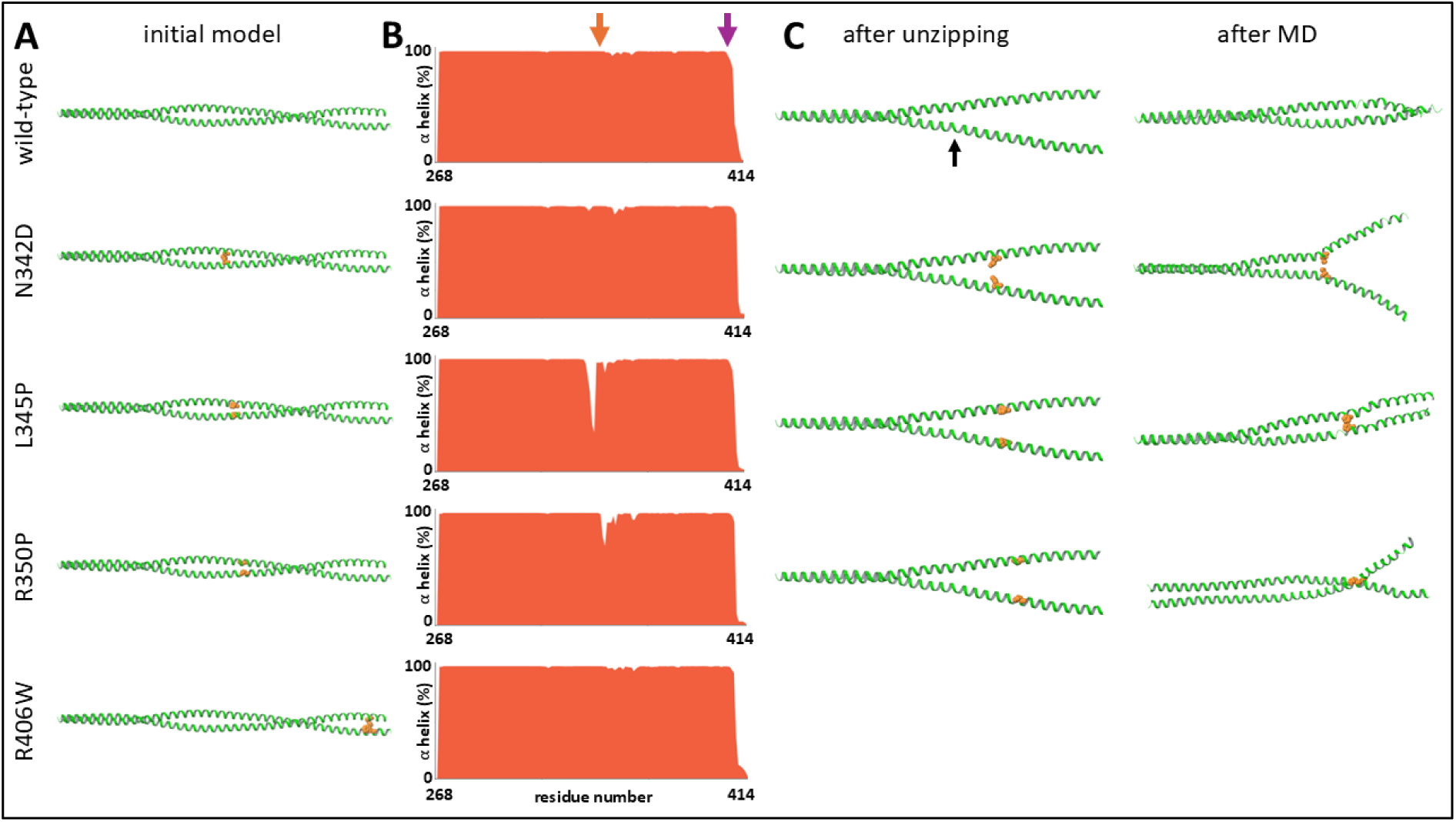
Molecular dynamics simulation of desmin coil 2 mutations. **(A)** Ribbon diagrams of the initial models with the mutated residues shown as orange balls. **(B)** Average α-helical propensity (%) over the course of a 100 ns MD simulation *versus* residue number. The arrows mark the position of center coil 2 mutations (orange) and R406W mutations (purple). **(C)** An MD simulation starting with the model unzipped at residue 328 (black arrow), and the conformation achieved after the simulation (5 ns for the wild-type, 10 ns for the mutants). The pronounced flexibility at the terminal region precludes a reliable assessment of coil 2 reassembly of the R406W mutant.

### Desmin mutant assembly

Next, we investigate the assembly kinetics of desmin mutants carrying the R406W, N342D, R350P, or L345P mutations. The time evolution of light scattering intensities at 405 and 594 nm for all four mutants differs markedly from that of wild-type desmin. All mutants except R406W show a significant delay before starting lateral assembly. Furthermore, once they start assembly, they exhibit a substantially stronger increase in scattering intensity compared to wild-type desmin (Figure 6, left column). The overall signal intensity of R406W-desmin is approximately 1.5-fold higher than that of wild-type desmin at 50 mM salt and about 3-fold higher at 100 mM salt. The characteristic shoulder that marks the transition from lateral to longitudinal assembly occurs at 900 ± 120 ms (50 mM) and 310 ± 50 ms (100 mM). Hence, the lateral assembly is slightly slower at 50 mM and slightly faster at 100 mM salt compared to wild-type desmin. The normalized intensity ratio I_594_/I_405_ after 10 min of assembly for both salt concentrations is increased compared to wild-type desmin and exceeds the Rayleigh-Gans limit I₅₉₄/I₄₀₅ = 1.46 for thin cylindrical rods, suggesting that R406W desmin does not form regular, elongated filaments.

**Figure 6.**
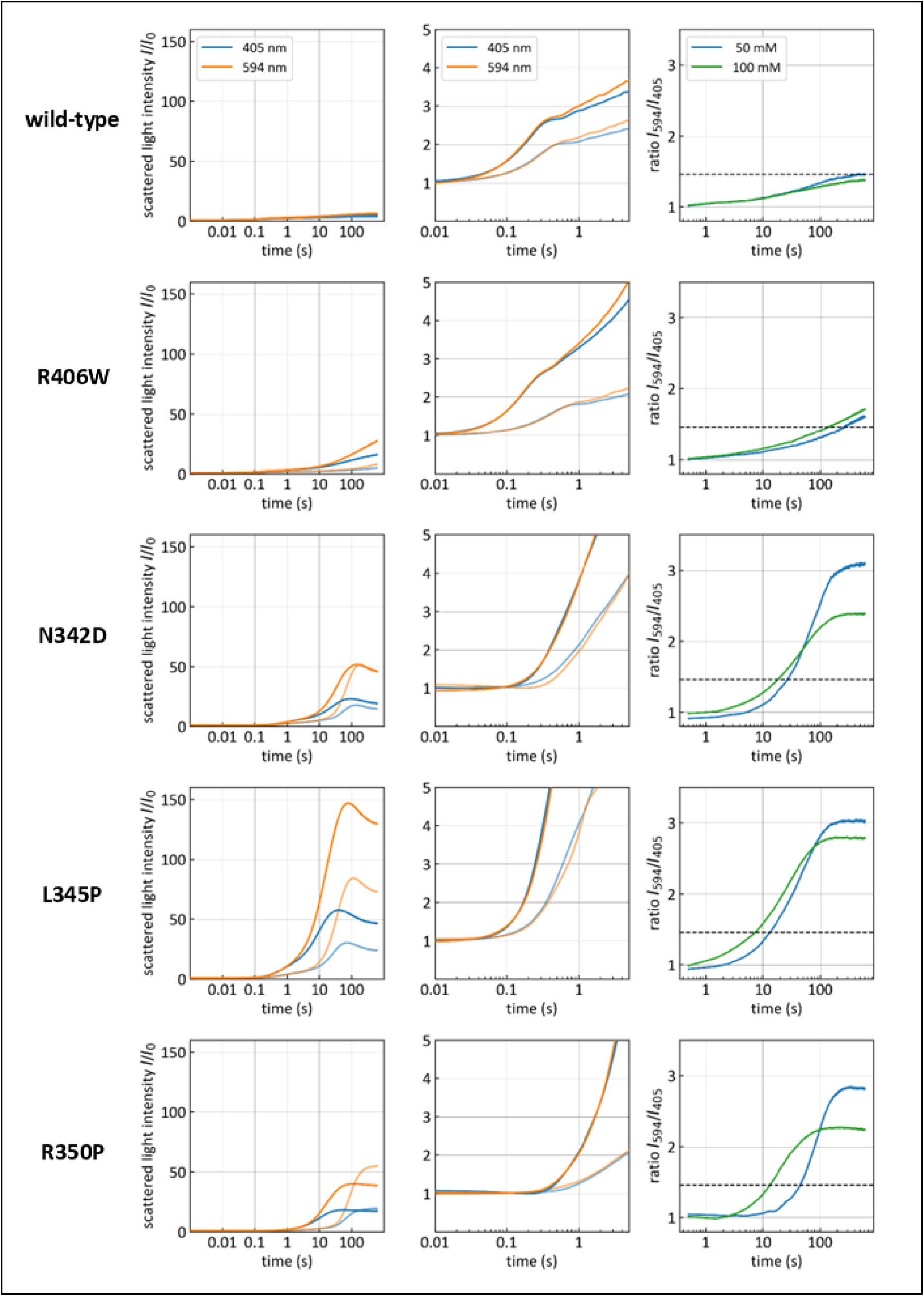
Assembly kinetics of wild-type desmin and desmin mutants measured with DWSF spectroscopy. Assembly of 0.1 mg/ml desmin is performed in 22.5 mM Tris-HCl buffer at 37°C at 50 mM (half transparent) or 100 mM (opaque) sodium chloride. Left column: Scattered light intensities at 594 nm (orange) and 405 nm (blue). The end point of the curves is at 600 s. Middle column: same data as shown in the left panels, zoomed in to the first 5 s. Right column: Scattered light intensity ratios I594/I405 at 50 mM (blue) and 100 mM NaCl (green). Dashed line shows theoretical maximal intensity ratio I594/I405 for wild-type desmin filaments, according to Rayleigh-Gans theory for thin cylindrical rods.

The desmin mutants N342D, L345P, and R350P produce scattering signals that are distinctly different from wild-type and R406W-desmin, yet similar to one another. For all three mutants, the scattering intensities reach levels between 2-fold and 10-fold higher than those of wild-type and R406W-desmin.

Strikingly, the onset of the signal increase is markedly delayed in all three mutants relative to wild-type and R406W-desmin (Figure 6; see also (19)). At 100mM salt the onset of signal increase is observed after approximately 150 ms for N342D-desmin and after 400 ms of R350P-desmin. At 50 mM salt, the corresponding onset times are approximately 300 ms and 600 ms, respectively, whereas wild-type desmin shows an increase already at about 30 ms, largely independent of salt concentration. Thus, relative to wild-type desmin, the onset of lateral assembly is delayed by approximately 5-fold and 10-fold for N342D, and by approximately 13-fold and 20-fold for R350P, at 100mM and 50mM salt, respectively. Although L345P exhibits only a comparatively small delay of signal increase, around 60 ms at 100 mM salt (2-fold) and around 100 ms at 50 mM salt (∼3-fold), it is still clearly delayed.

These delay times of the lateral assembly indicate that these mutations stay unusually long in a tetrameric conformation after salt addition. Furthermore, at 50 mM salt, the effect of the mutation on the lateral assembly is more pronounced than at 100 mM, as reflected by the larger fold-delays relative to wild-type and R406W-desmin. However, once assembly begins, these mutants exhibit a substantially greater increase in signal intensity than wild-type and R406W-desmin. Moreover, none of the three mutants displays the characteristic shoulder associated with the transition from lateral to longitudinal assembly. Together, these observations identify delayed lateral assembly followed by an abrupt increase in signal intensity as a prominent shared phenotype of the coil 2 center mutants, a behavior not observed for R406W desmin.

Due to this altered behavior during the lateral assembly phase, we cannot determine from the absolute signal intensity alone, if the seemingly higher rate of the signal increase at the later time points reflects faster kinetics or a larger scattering cross section of the assembly products. However, we find that the intensity ratios I_594_/I_405_ of N342D, L345P and R350P-desmin reach considerably higher values than those of wild-type and R406W-desmin (Figure 6, right column). After approximately 10 s of assembly, the ratio exceeds the Rayleigh-Gans limit I_594_/I_405_ = 1.46 for thin cylindrical rods. This suggests that none of the three mutants in the center of coil 2 form elongated filaments beyond 10 s. Instead, we observe a mixture of filaments and an increasing number of aggregated assembly products. Due to their larger scattering cross-section, the aggregates are primarily responsible for the pronounced increase in signal intensity.

**Table 1.** Shape and size of the assembly products of R406W, N342D, L345P, and R350P-desmin. The assembly is performed at a protein concentration of 0.1 mg/ml at 100 mM salt for 1 s and 5 min (for L345P for 3 min) at 37°C. Corresponding AFM images are displayed in Figure 7. ULF: unit-length filaments; SF: short filaments.

|  | **R406W** | **N342D** | **L345P** | **R350P** |
| --- | --- | --- | --- | --- |
| **1 s** | ULFs, SF, some clusters | Clusters of SF (~300 nm) | Mix of SF and globular assemblies ($\phi \sim 100\text{-}200$ nm) | Mix of ULFs, SF and globular assemblies ($\phi \sim 100\text{-}200$ nm) |
| **5 min** | Large clusters (~10 $\mu\text{m}$) of SF | Aggregates ($\phi \sim 1$ $\mu\text{m}$) of small globular assemblies | Clusters ($\phi$ 1-2 $\mu\text{m}$) of globular assemblies ($\phi \sim 100\text{-}200$ nm) | Clusters ($\phi$ 1-2 $\mu\text{m}$) of globular assemblies ($\phi \sim 100\text{-}200$ nm) |

To visualize the assembly products, we image them at the beginning (1 s) and the actual end (5 min) of the assembly process in 100 mM salt using AFM (Figure 7). The shape and approximate size of the assembly products are summarized in Table 1. For all four mutants, we find that the assembly products strongly differ from the typical single elongated filaments of wild-type desmin (Figure 3). Moreover, we find for all four mutants a hierarchical structure of the assembly products, consisting of clustered short filaments or globular complexes. These globular complexes have a size of 100-200 nm and are visible already after 1 s of assembly. Interestingly, the N342D, L345P and R350P mutant initially (after 1 s) assemble also into short filaments (1 - 3 ULFs), in parallel with globular complexes, but by 1 min, filaments have disappeared and only clusters of globular complexes remain.

**Figure 7.**
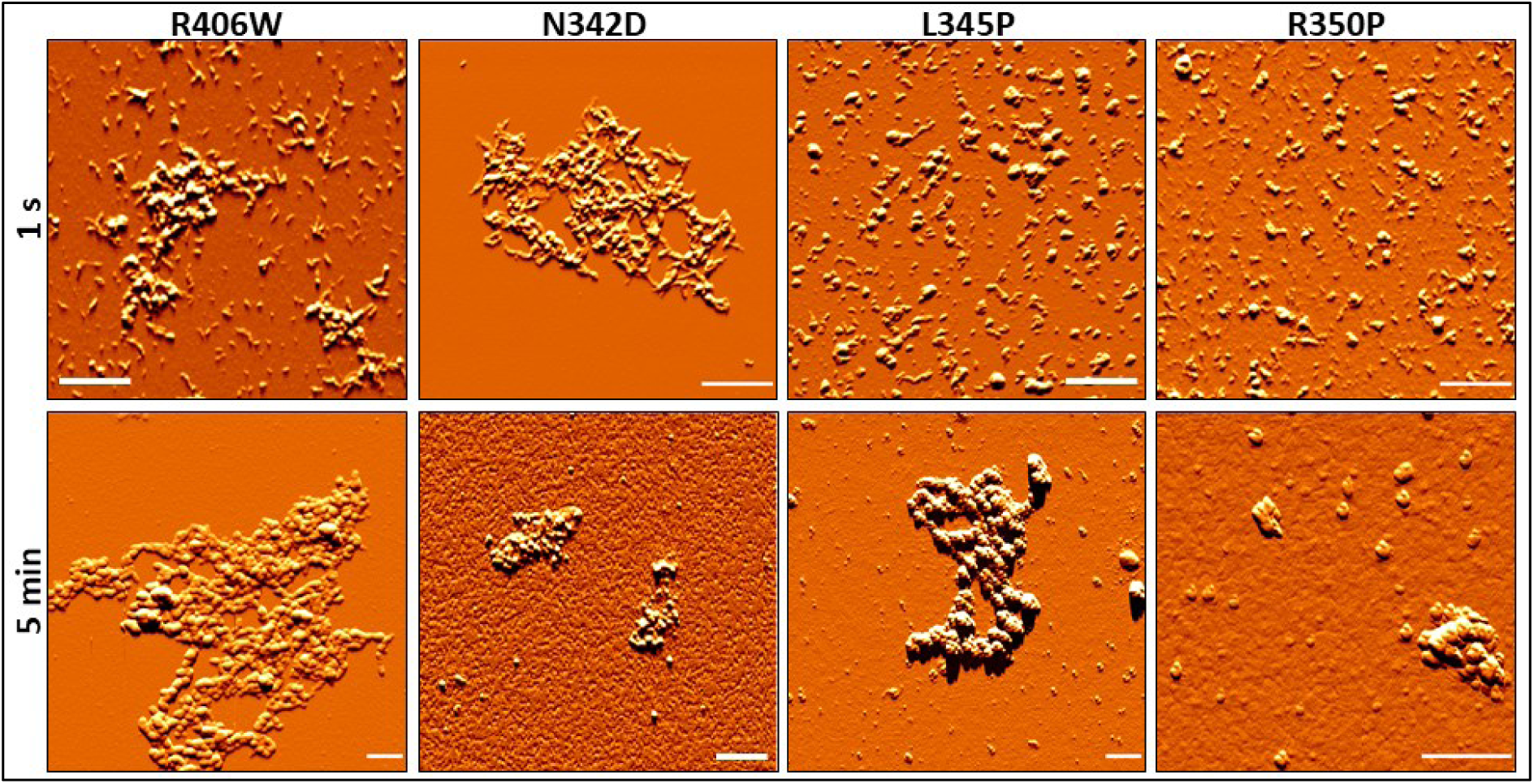
Atomic force microscopy of mutant desmin assembly. Proteins are assembled at a concentration of 0.1 mg/ml in 22.5 mM Tris-HCl buffer, pH 7.5, containing 100 mM salt at 37°C for 1s and 5 min (for L345P we show an image taken at 3 min instead of 5 min). Scale bar: 1 µm.

### Assembly of equimolar mixtures of mutant and wild-type desmin

The desmin mutants investigated here arise in humans from autosomal dominant mutations and are therefore heterozygous. Because both genes are expressed at the same level, both wild-type and mutant proteins are present in roughly equal amounts. A previous study has reported that the presence of equal amounts of wild-type and mutant desmin partially rescues assembly (51). We therefore tested whether this also holds for the L345P mutant and compared its assembly with the assembly R406W mutant using DWSF. We focus in the following on R406W-desmin, which retains the ability to form fibrous structures, and L345P-desmin, which predominantly forms globular structures. AFM images taken after 1 and 5 minutes of assembly of wild-type and R406W-desmin mixtures reveal a similar behavior as pure R406W mutants: Filaments remain short (316 ± 44 nm, ∼ 7 ULFs) after 5 min of assembly and are predominantly found in carpet-like clusters, although these appear less densely packed compared to pure mutants. Apparently, the presence of wild-type desmin reduces lateral adhesiveness by forming mixed filaments and thus partly rescues filament formation (Figure 8 A, B).

For equimolar wild-type and L345P-desmin mixtures, we observe similar small globular structures (140 ± 31 nm and ∼3 ULFs), as obtained for the mutant alone after 1 and 5 minutes. However, these globular structures exhibit a less pronounced propensity to form large clusters, mirroring the tendency observed in the wild-type and R406W-desmin mixtures. Unlike R406W-desmin, mixing L345P-desmin with wild-type desmin does not offset the mutant’s destructive impact on filament formation, as no filaments are detected (Figure 8 C, D).

**Figure 8.**
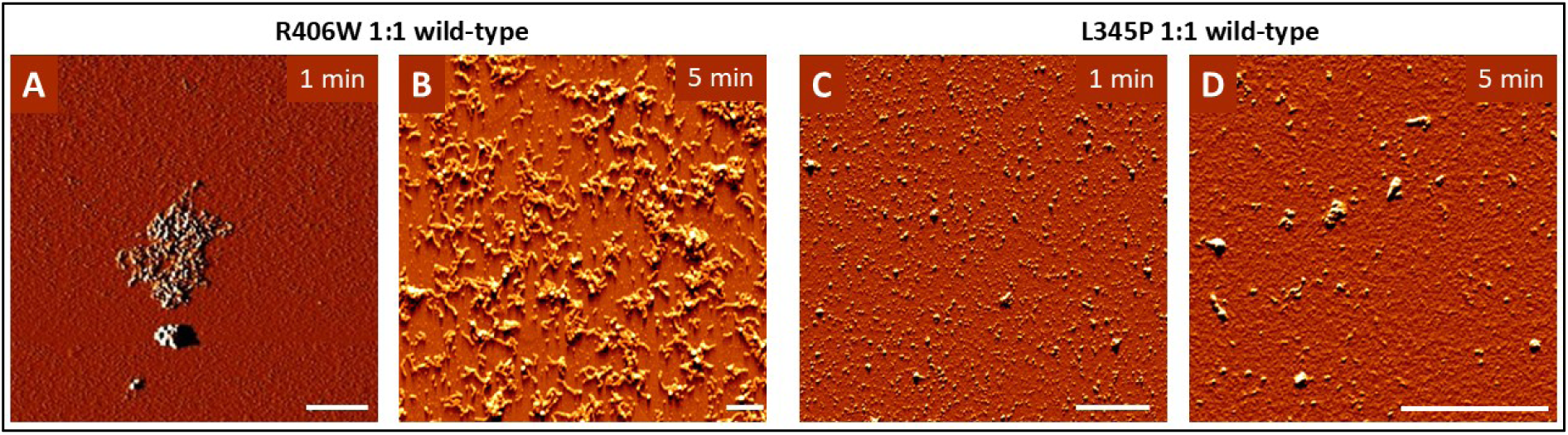
Atomic force microscopy of R406W-wild-type desmin mixture (equal amounts; A, B), and L345P-wild-type desmin mixture (equal amounts; C, D). Proteins are assembled at a concentration of 0.1 mg/ml in 22.5 mM Tris-HCl buffer, pH 7.5, containing 100 mM salt at 37°C for 1 min and 5 min. Scale bars: 2 µm.

**Figure 9.**
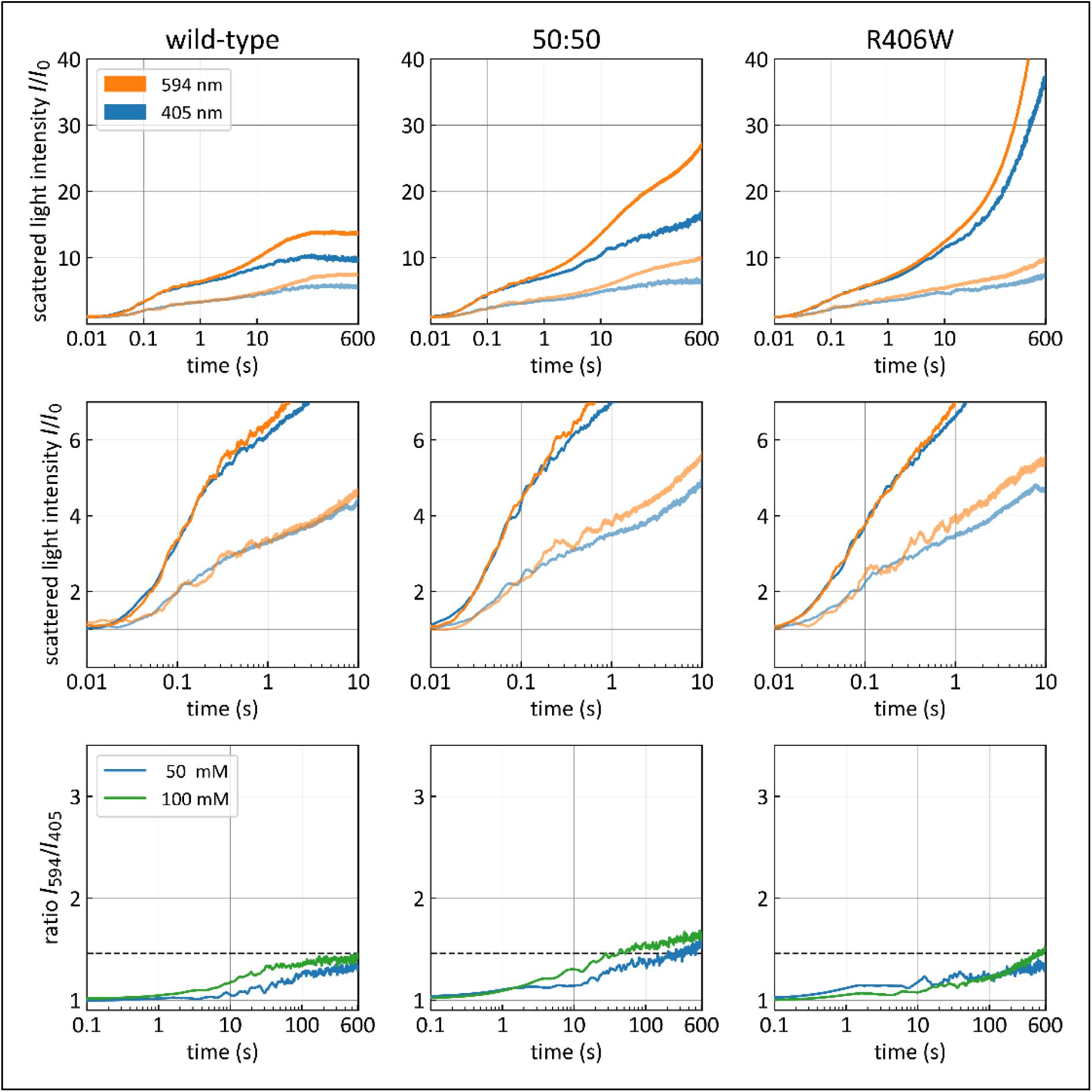
DWSF assembly profiles of wild-type desmin (left panels), equimolar mixtures of wild-type and R406W-desmin (middles panels), and R406W-desmin (right panels). Proteins are assembled at a concentration of 0.1 mg/ml in 2 mM Mops, pH 7.5, with 50 mM (half transparent) or 100 mM (opaque) potassium chloride, at 37 °C. Upper row: light scattering intensities recorded over 600 s at 594 nm (orange) and 405 nm (blue). Middle row: zoom-in of upper row to 0.01 s to 10 s. Lower row: intensity ratios I594/I405. The dashed lines indicate the Rayleigh-Gans limit for thin cylindrical rods of 1.46.

DWSF measurements of one-to-one mixtures of wild-type and R406W-desmin also reveal behavior that falls between that of pure wild-type and pure R406W-desmin. (Figure 9). First, during the lateral assembly phase, the scattering signal closely resembles that of wild-type desmin, including the pronounced inflection point. Subsequently, the signal rises somewhat more steeply for the mixture than wild-type desmin but does not reach the intensity of R406W-desmin alone. The 594 nm and 405 nm light signals for the R406W mixtures diverge at both salt concentrations, but later than in wild-type assembly. After about 20 s of assembly, the intensity ratio I_594_/I_405_ of the mixture exceeds that of wild-type desmin and exceeds the Rayleigh-Gans limit for thin cylindrical rods of 1.46. This suggests that the mixture forms both wild-type-like filaments and lateral aggregates of filaments.

**Figure 10:**
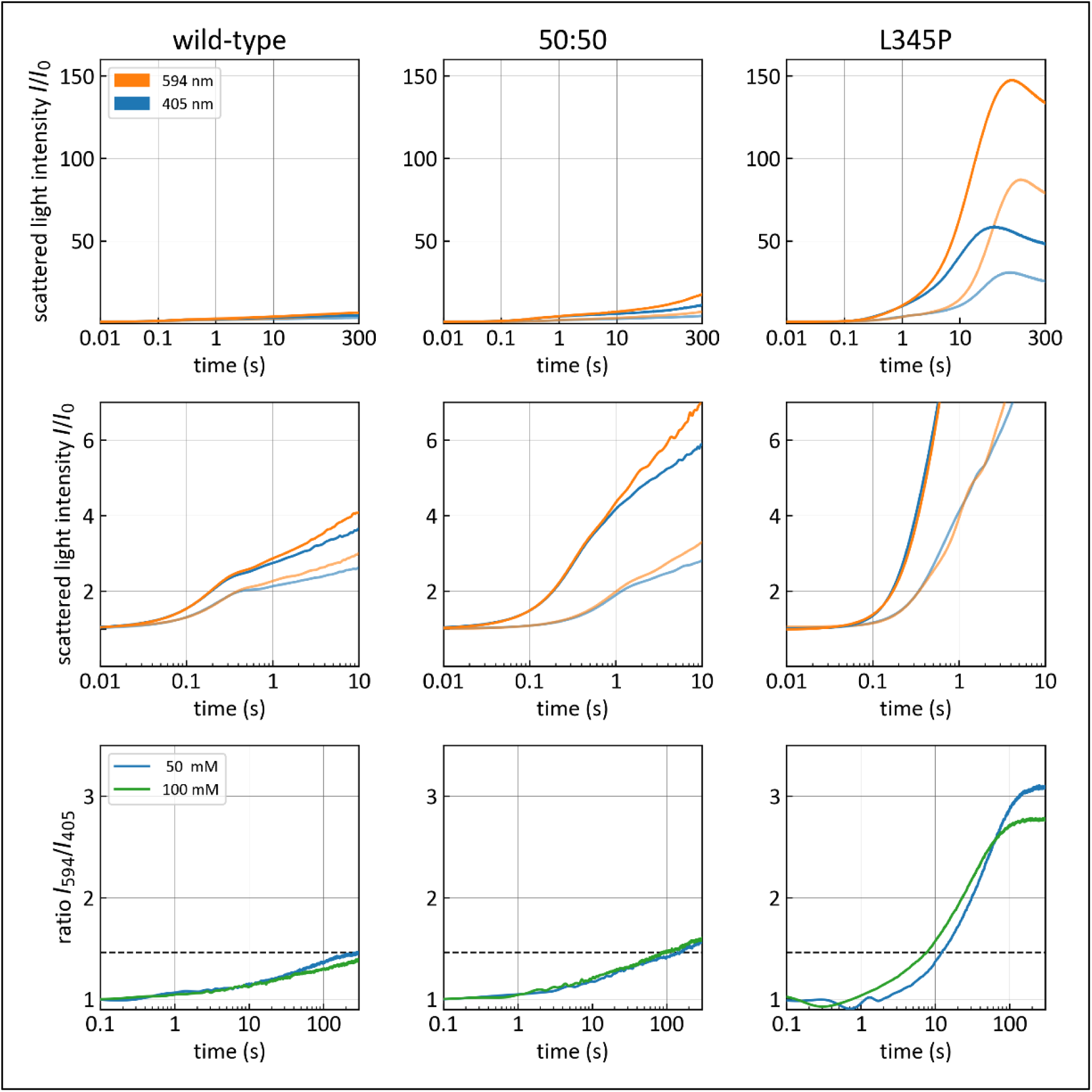
DWSF assembly profiles of wild-type desmin (left panels), equimolar mixtures of wild-type and L345P-desmin (middles panels), and L345P-desmin (right panels). Proteins are assembled at a concentration of 0.1 mg/ml in 2 mM Mops, pH 7.5, with 50 mM (half transparent) or 100 mM (opaque) potassium chloride, at 37 °C. Upper row: light scattering intensities recorded over 300 s at 594 nm (orange) and 405 nm (blue). Middle row: zoom-in of upper row to 0.01 s to 10 s. Lower row: intensity ratios I594/I405. The dashed lines indicate the Rayleigh-Gans limit for thin cylindrical rods of 1.46.

DWSF assembly profiles of equimolar mixtures of wild-type and L345P-desmin also fall in between that of pure wild-type and mutated desmin (Figure 10). The mixtures display a shoulder at the transition from lateral to longitudinal assembly, albeit less pronounced compared to pure wild-type desmin. Subsequently, the signal increases faster than for pure wild-type desmin but much slower than for mutated desmin. Furthermore, the intensity ratio I_594_/I_405_ of the protein mixture only slightly exceeds the Rayleigh-Gans limit of 1.46. These observations confirm the conclusions drawn from our AFM measurements, namely that presence of wild-type desmin partially rescues the uncontrolled aggregation of L345P-desmin and reduces the formation of large protein clusters.

### Numerical simulation of the light scattering signals of desmin mutants

All four mutants assemble into distinct types of short fibrous or globular structures. Consequently, their light scattering behavior differs from that of isolated elongated wild-type desmin filaments, which is well described by the Rayleigh-Gans theory for cylindrical rods. This theory predicts that after the filaments elongate beyond a length of 6-7 ULFs (∼300 nm), the signal intensity ratios at 594 nm and 405 nm cannot increase beyond a value of 1.46 (49, 50). Experimentally, we find that the intensity ratios I_594_/I_405_ of the mutant proteins exceed this limit, and hence the Rayleigh-Gans regime for cylindrical rods does not apply. We therefore explore whether information about the shape and size of the assembly products can be gained by analyzing the intensity ratio I_594_/I_405_. To do so, we turn to numerical simulation of light scattering.

Previous studies have explored the light-scattering of worm-like chain (WLC) protein aggregates, and protein aggregates with a fractal dimension below 3 (54, 56, 73). Both may be relevant in the context of the present study: Short R406W-desmin filament bundles exhibit curved, random-coil-like conformations, consistent with worm-like chains of low apparent persistence length (Figure 7, (51)). The other desmin mutants arrange in the shape of differently packed globules resembling fractal structures.

To simulate the light scattering of protein aggregates, we consider them to consist of chains of spheres with a diameter of 12.5 nm, in the following referred to as “beads”. We choose this value because it corresponds to the diameter of desmin filaments. For example, a single ULF could then be modeled as a linear chain of 5 beads.

The WLC filament length is varied by adjusting the number of connected beads. Each bead can bind only to two neighboring beads. The internal conformational statistics of this chain of beads is described using the worm-like chain model, and the light scattering signal from each bead as well as the light interference between all beads within a chain is computed using Debye scattering theory, as described in Methods. Scattered light from neighboring WLCs is assumed not to interfere with each other.

First, we use this model to reproduce the experimentally measured scattering behavior of wild-type desmin. Reported persistence lengths of wild-type desmin filaments range between 300 nm and 1000 nm (59). Within this range, the simulated intensity ratio shows little dependence on the persistence length (Figure 11 A). For WLC with a contour length up to one ULF (∼ 60 nm), the intensity ratio I_594_/I_405_ remains near unity. With increasing contour length, the ratio value rises until it saturates at around 1.46 for lengths above 300 nm length (Figure 11, dashed line). This behavior perfectly agrees with the predictions by the Rayleigh-Gans theory for straight rods (Figure S6) and with our experimental data of wild-type desmin: the ratio increases during the onset of longitudinal assembly and saturates near 1.4 as the filament length approaches the length of the scattering wavelength (Figure 11 B).

**Figure 11.**
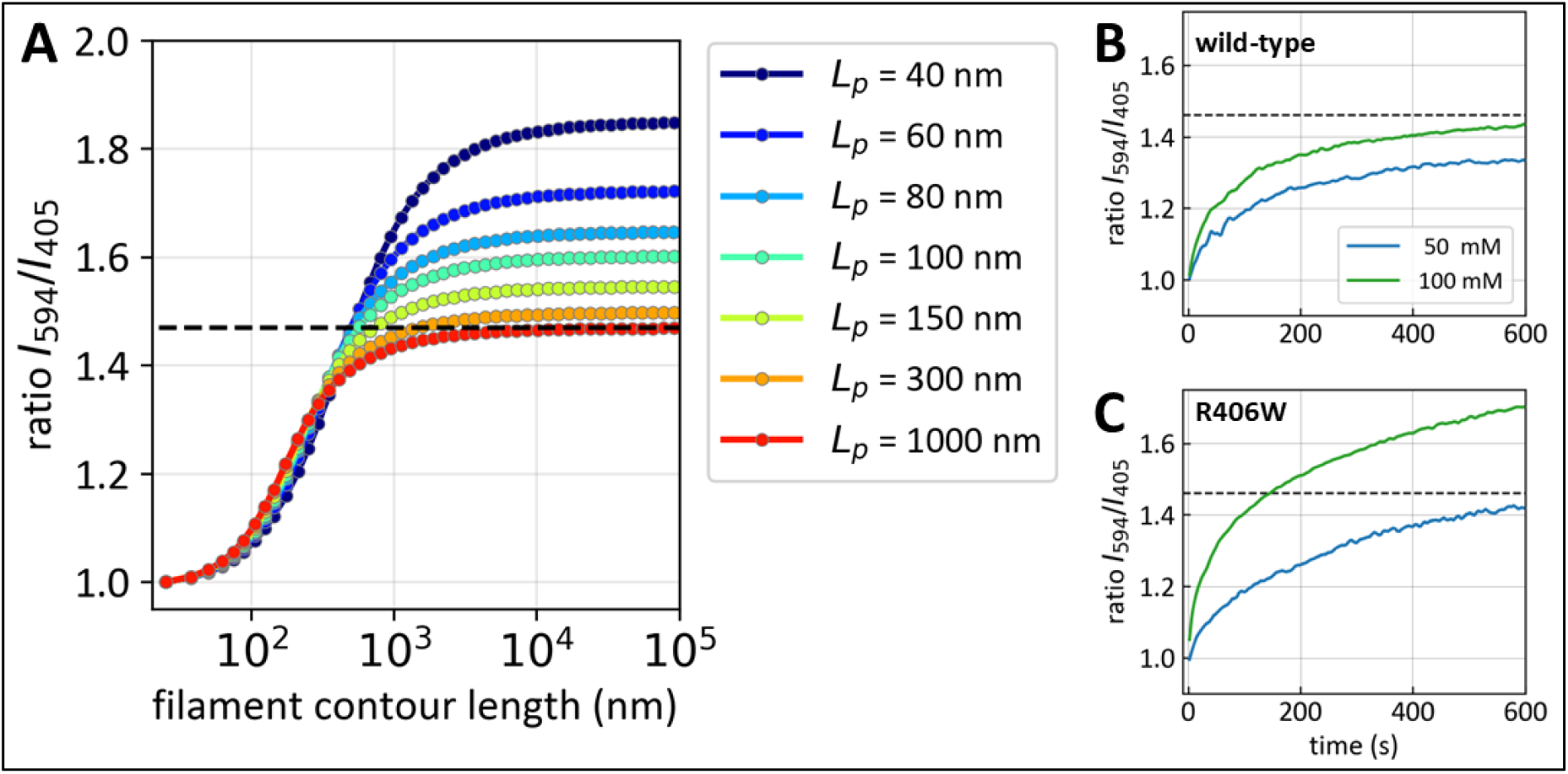
Simulated scattered light intensity ratios using the WLC model. **(A)** Simulated scattered light intensity ratio I594/I405 over contour length of filaments approximated as a worm-like chain. The colors depict different persistence length from *L_p_*=1000 nm down to 40 nm. Filaments with contour lengths ranging from 25 nm to 100 µm are sampled from a logarithmically uniform distribution. Measured example intensity ratios of wild-type desmin **(B)** and R406W-desmin **(C)** at 50 mM (blue) and 100 mM salt (green). The dashed lines indicate the Rayleigh-Gans limit for thin cylindrical rods of 1.46.

R406W-desmin clusters have the appearance random coils of WLCs. These clusters exhibit a pore size that roughly corresponds to the persistence length of a WLC. We do not suggest that R406W-desmin aggregates form through the bending, curling and entanglement of longer filaments. Rather, the shape of WLCs serves as a geometrically similar structure with a similar scattering behavior (Figure 11, Figure 13). We find that the scattering behavior of a WLC with a persistence length of 100 nm best recapitulates the intensity ratio I_594_/I_405_ of R406W-desmin clusters after 5 min of assembly (Figure 11 A, C). This reduced persistence length indicates a weaker inter-ULF repulsion compared to wild-type desmin. This agrees with the apparent spacing (or pore size) between short rod-like assembly products observed with AFM (Figure 7, after 5 min).

At 50 mM salt, DWSF spectroscopy measurements exhibit an intensity ratio of 1.3 ± 0.1 after 10 min of assembly, suggesting that the assembly of long filaments is impaired under low salt conditions and does not continue beyond 300 nm or 6 ULFs. EM and AFM images confirm a stop of assembly after a few ULFs (13), which furthermore implies that the scattering behavior is dominated by Rayleigh scattering and not influence by the persistence length.

In one-to-one mixtures of wild-type desmin with R406W-desmin, the I_594_/I_405_ plateau values remain increased, and the scattering signal is dominated by short persistence length WLC-like assemblies, despite the observation with AFM (Figure 8) that some isolated, non-aggregated filaments are present after 5 min of assembly.

**Table 2.** Averaged results DWSF spectroscopy measurements for 0.1 mg/ml wild-type and mutant desmin at 50 mM and 100 mM salt. Kinetics is measured as negative exponential fit to the intensity ratio I594/I405 (mean ± SD) with open fit to the ratio plateau (saturation level). The ratio plateau is given as an average of 5-10 different measurements for each measurement condition (mean ± SD). The fractal dimension is derived from the average plateau values.

| DESMIN | Time constant of<br>exponential fit to ratio [s] | | Ratio Plateau<br>$I_{594}/I_{405}$ | | Fractal Dimension | |
| --- | --- | --- | --- | --- | --- | --- |
|  | 50 mM | 100 mM | 50 mM | 100 mM | 50 mM | 100 mM |
| WILD-TYPE | 76 ± 34 | 57 ± 22 | 1.28 ± 0.13 | 1.41 ± 0.15 | 1 | 1 |
| R406W | 141 ± 133 | 107 ± 107 | 1.27 ± 0.13 | 1.57 ± 0.14 | 1 | 1.2 ± 0.2 |
| N342D | 133 ± 85 | 102 ± 59 | 3.16 ± 0.19 | 2.55 ± 0.49 | 3 ± 0.2 | 2.4 ± 0.5 |
| L345P | 78 ± 45 | 51 ± 27 | 2.79 ± 0.63 | 2.66 ± 0.37 | 2.7 ± 0.6 | 2.6 ± 0.4 |
| R350P | 83 ± 25 | 29 ± 5 | 2.42 ± 1.63 | 2.77 ± 0.53 | 2.3 ± 1.7 | 2.7 ± 0.5 |
| VIMENTIN | 258 ± 52 | 120 ± 18 | n.d. | n.d. | 1 | 1 |

An alternative approach to model the aggregation behavior of R406W-desmin is the fractal dimension analysis. This is a standard approach to describe the aggregation reaction of colloidal systems (54, 55). There, a diffusion-limited aggregation reaction results in fractal dimensions of around 1.7, whereas reaction-limited aggregation results in fractal dimensions of around 2.0-2.2 (74). As described below, different fractal dimensions result in specific intensity ratios. For the intensity ratio of 1.6 that we measure for the R406W variant, the corresponding fractal dimension is 1.2. The visualization of R406W protein aggregates with that fractal dimension (Figure 13 B) closely agrees with AFM measurements (Figure 7).

Fractal dimension analysis becomes important for analyzing the scattering behavior of the coil 2-center mutants N342D, L345P, and R350P, which form spherical protein clusters and not random coils, as shown by AFM and previously by EM (Figure 7; (13)). Accordingly, they exhibit intensity ratio I_594_/I_405_ plateau values between 2.4 and 2.8 (Table 2). These values are beyond the range that can be predicted with the WLC model. To explain the scattering behavior of the center coil 2 mutants, we describe their packing arrangements and densities by the fractal dimension D_f_, which provides a model-dependent estimate to describe the mass distribution within a cluster. The fractal dimension spans the range 1 ≤ D_f_ ≤ 3, corresponding to structures ranging from open, chain-like assemblies (D_f_ ≈ 1) to compact, densely packed (D_f_ → 3) spheres.

The simulated intensity ratio I_594_/I_405_ monotonically increases with cluster radius up to 250-300 nm and then saturates, depending on the fractal dimension (Figure 12 A). At a fractal dimension of unity, the simulation slightly underestimates the prediction from the Rayleigh-Gans theory, because at the single element level (ULF), the size of the scattering object is already slightly beyond the Rayleigh limit.

**Figure 12.**
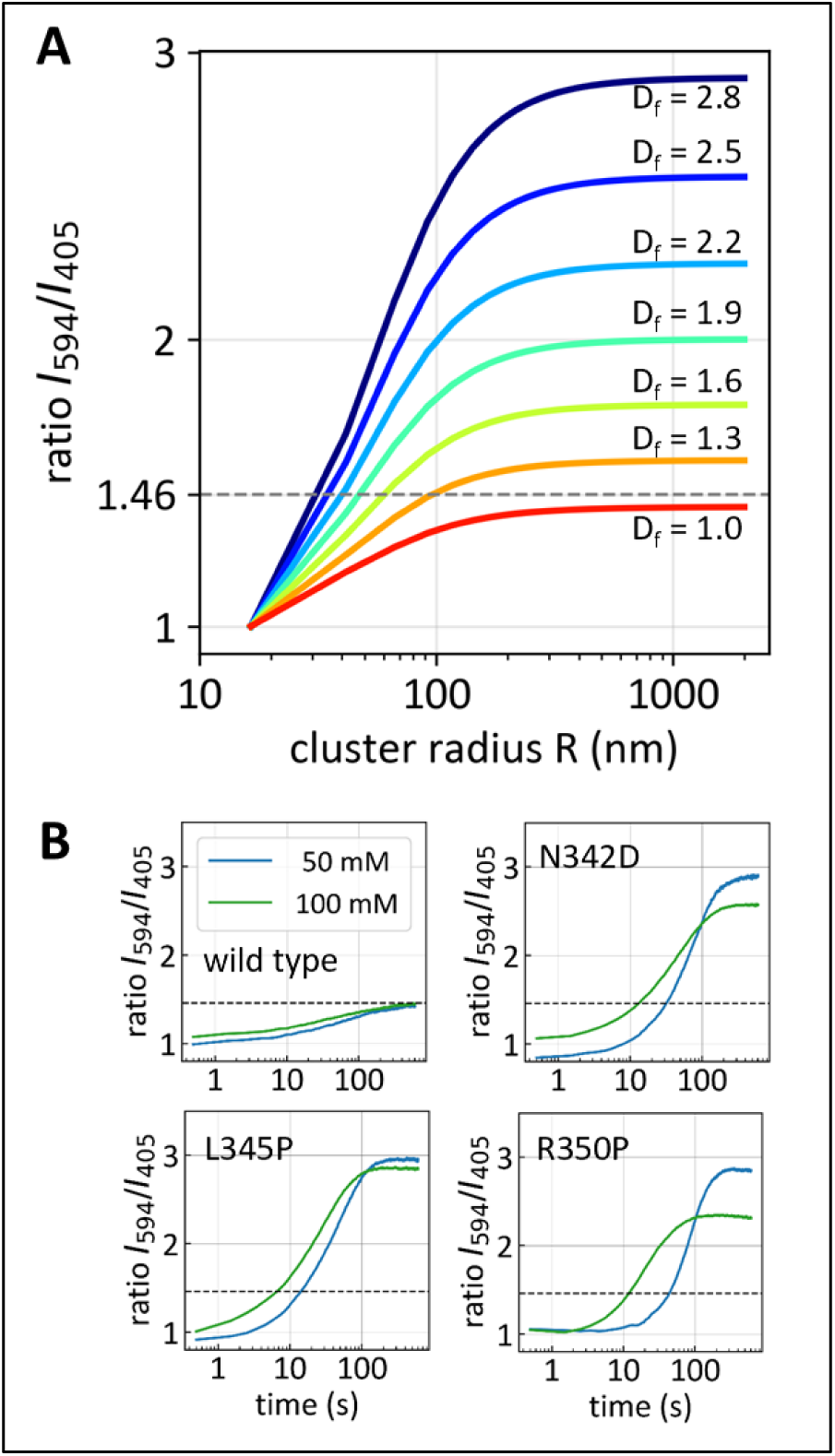
Simulated scattered light intensity ratios using the fractal dimension model. **(A)** Simulated scattered light intensity ratio I_594_/I_405_ over increasing aggregation cluster radius for different fractal dimensions. The fractal dimensions depict the internal structure of each cluster from loosely filamentous (D_f_ = 1, red) with a saturation below a ratio of 1.46 to dense, near solid-like packing (D_f_ = 2.8, dark blue). **(B)** Measured intensity ratios for wild-type, N342D-, L345P-, R350P-desmin at 50 mM (blue) and 100 mM salt (green).

Using the straightforward conversion between the plateau ratio and the fractal dimension (Eq. 12, Figure 12 A), we find that the fractal dimensions of the N342D, L345P, and R350P variants fall between 2.3 and 2.7, for both salt concentrations (Table 2). Such high fractal dimension values suggest a much more compact mass distribution than filaments have, which is consistent with the clustered morphologies observed in the AFM images (Figure 7). Only N342D-desmin aggregates at 50 mM salt show on average an intensity ratio I_594_/I_405_ above three and are therefore outside of this regime. EM images of this protein at 50 mM salt showed sheets of granulofilamentous material (15). Therefore, they are not covered by the fractal dimension model. One-to-one mixtures of wild-type desmin and L345P mutant proteins exhibit smaller fractal dimensions of around 1.2.

We use the two presented models, WLC and fractal dimension analysis, to simulate protein aggregate morphologies that can be directly compared to our AFM images. In the worm-like chain representation, wild-type desmin with a persistence length of 1000 nm forms extended, only weakly curved filaments that span the simulation box (Figure 13 A) and resemble the long, continuous filaments observed by AFM (Figure 3). By contrast, R406W-desmin is described by much shorter and more flexible chains with a persistence length of 100 nm, which collapse into compact, entangled filament bundles in projection (Figure 13 A), consistent with the shorter and more disordered filamentous structures in the AFM data (Figure 7).

The fractal aggregation simulations translate the same scattering constraints into assemblies of 40 nm spheres, corresponding to the globular complexes apparent in the AFM images (Figure 7). For R406W-desmin, a low fractal dimension of 1.2 yields open clusters of loose patches of aggregates (Figure 13 B), as seen experimentally. By contrast, the N342D, L345P, and R350P mutants are reproduced only by dense globular clusters with a fractal dimension of around 2.5, which generate small, compact aggregates (Figure 13 B), matching the tightly packed globular structures in the AFM images (Figure 7). Together, these simulations establish a connection between DWSF light scattering-derived structural parameters and the morphology seen in AFM and EM images. Our simulations furthermore demonstrate that the scattering behavior at an intermediate range of intensity ratios I_594_/I_405_ below 1.8 can result from either flexible filament bundles or open globular clusters. However, the more strongly scattering mutants are better explained by compact globular aggregates than by filamentous assemblies.

**Figure 13.**
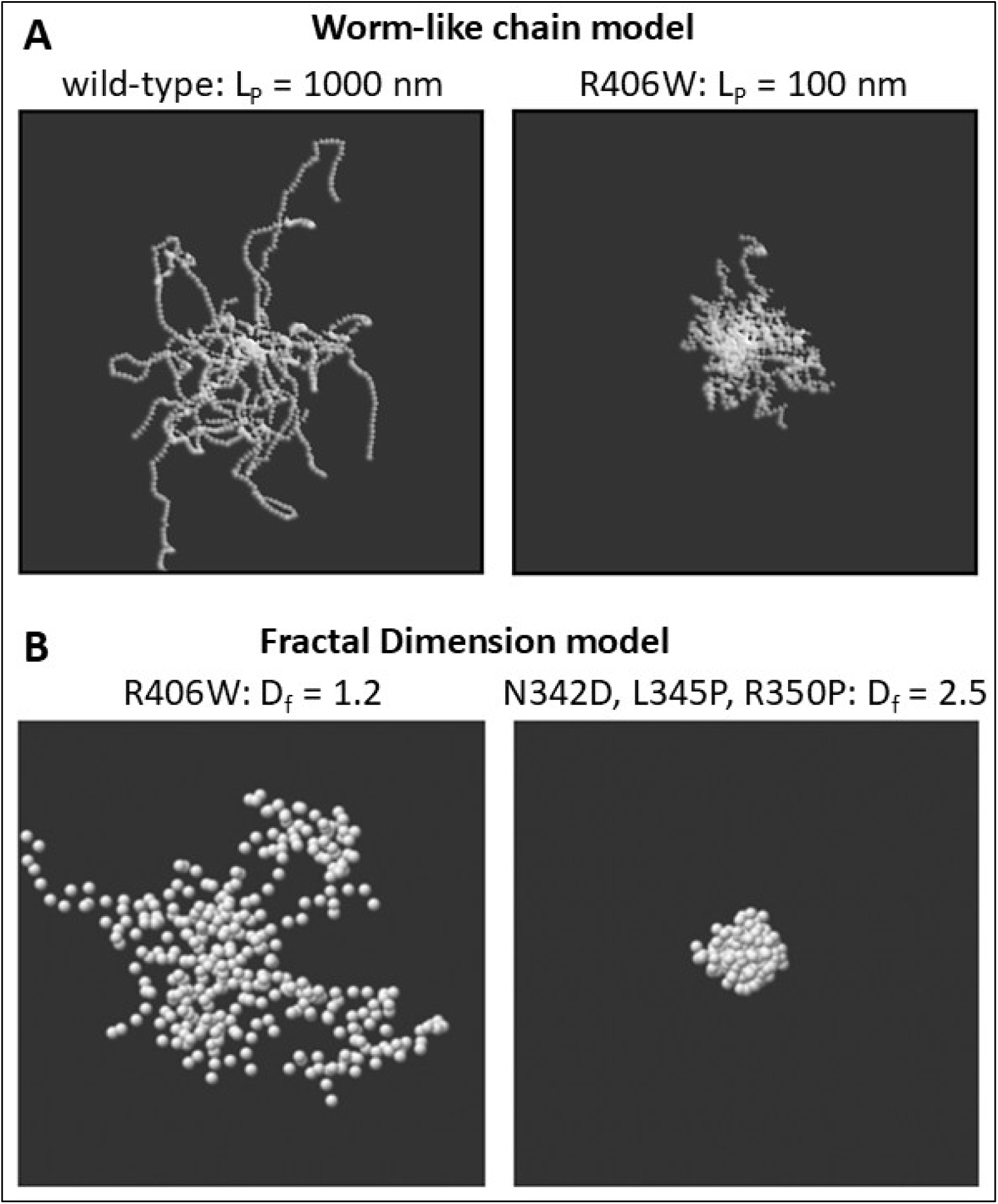
Topological simulation of protein aggregates in a two-dimensional projection. **(A)** Worm-like chain model with a persistence length LP = 1000 nm corresponding to wild-type desmin filaments (left) and LP = 100 nm for R406W-desmin (right). **(B)** Globular particles seeded according to the fractal dimension model. Fractal dimension Df = 1.2 for R406W-desmin aggregates (left); Df = 2.5, representing N342D, L345P, R350P-desmin aggregates (right). Simulations are seeded around center point and consist of 20 WLCs or 500 globular particles per cluster.

## DISCUSSION

A central finding of this study is that the in vitro assembly of all four desmin mutants is defective from the earliest detectable stage. Abnormalities arise within the first 300 milliseconds after salt is added, as determined by DWSF spectroscopy with millisecond time resolution. The formation of the first higher-order complexes from tetrameric subunits (i.e., octamers) is significantly delayed after salt addition compared to wild-type desmin assembly. This delay depends on both the specific mutation and the salt concentration. Nevertheless, AFM observations suggest that the primary assembly process of ULFs is comparable for mutant and wild-type desmin. However, once the ULFs anneal into short filaments during the early phase of assembly, productive elongation of mutant assembly abruptly ceases. Instead, short filaments formed by the N342D, L345P, and R350P mutants appear to reorganize segment by segment into small globular complexes that remain connected (15). In contrast, the R406W mutant forms micrometer-long bundles of laterally associated filaments that subsequently aggregate into even larger complexes.

These mutation-specific differences in lateral assembly behavior are most likely determined by the chemical nature of the substituted residue and position within the heptad and hendecad repeat patterns of the coiled coil (Figure 1 A ii). To investigate this possibility, we performed MD simulations of coiled-coil dimers encompassing the entire coil 2 domain. All three center-of-coil-2 mutations produce pronounced abnormalities in structural dynamics, indicating that these substitutions perturb the native conformational landscape of the dimer.

Such dynamic defects may have direct consequences for ULF assembly. They may expose interaction surfaces that are inaccessible in the wild-type state or alter tetramer geometry prior to ULF formation. Consistent with the latter possibility, analytical ultracentrifugation experiments in low-salt conditions revealed lower sedimentation coefficients for the two coil 2 proline mutants (4.8 S for R350P and 4.9 S for L345P), compared to the normal wild-type desmin tetramers (5.2 S) (15). These reduced values suggest that the proline substitutions induce conformational loosening, increased flexibility, or altered dimer-dimer packing prior to salt-induced assembly. In contrast, R406W-desmin exhibits a slightly higher *s*-value of 5.4 S, which is consistent with a more compact tetrameric state. However, alternative changes in shape or hydration cannot be ruled out (15, 75). The markedly delayed onset of lateral assembly observed for N342D and R350P is consistent with such structural perturbations.

The comparatively modest delay observed for L345P, in turn, correlates with the relatively rapid recovery of locally unzipped coiled coils in the MD simulations. The situation is different for R406W-desmin. MD simulations are less informative for this mutation because the terminal region of coil 2 is intrinsically flexible. Nevertheless, the high conservation of the final 30 amino acids of coil 2 suggests that replacing arginine with tryptophan affects assembly. Indeed, R406W-desmin exhibits wild-type-like kinetics during the initial 30 seconds of assembly (51). This indicates that ULF formation and initial filament elongation are largely preserved. The defect emerges later, when lateral filament-filament interactions become increasingly dominant and lead to the formation of filament bundles that impede the elongation of individual filaments.

These longitudinal assembly defects raise a central mechanistic question: How are interactions between two ULFs initiated, and how do mutations in the central coil domain affect this process? One possibility is that productive interaction begins with head-to-tail contact between dimers from neighboring ULFs (brackets in Figure 1 B iii). Alternatively, the initial step could involve a weak, dynamic interaction between antiparallel coil 2 segments (Figure 1 B iii), guided by the spatial pattern of ionic clusters along coil 2 (76). The experiments reported here indicate that longitudinal assembly is mostly guided by coil 2–coil 2 interactions, and less so by head-to-tail interactions. The R406W mutation, located in the head-to-tail overlap region, has no measurable effect on early filament elongation. In contrast, the longitudinally defective central coil 2 mutations may affect the coil 2–coil 2 interactions. Previous evidence supporting the importance of coil 2–coil 2 interactions was through a mouse desmin mutant (1–402), which lacks in addition to the entire tail segment a part of the consensus motif of coil 2 . Even though this mutant was expected to be compromised in the head-to-tail interaction, it retains the ability to undergo longitudinal elongation, though the radial compaction is delayed (77).

Replacing arginine with tryptophan may alter intermolecular interactions, diverting assembly from its normal pathway. Substituting the strongly native basic arginine with tryptophan increases the local hydrophobicity of the dimer surface and disrupts an interhelical salt bridge with Glu401. At the same time, the polarizable π-electron system of the indole ring creates a markedly different electrostatic and interaction environment (78). Together, these changes are likely to shift the balance between specific and nonspecific intermolecular interactions, thereby generating adhesive filament surfaces. Thus, R406W does not primarily impair the initial stages of assembly but instead promotes aberrant lateral association between already formed filaments.

A hallmark of normal desmin assembly is the progressive reduction in apparent filament diameter during maturation. This radial compaction occurs between 2 seconds to 10 minutes and reflects an essential structural transition from initially loose filament precursors to mature, compact IFs (27, 32). In wild-type desmin, this process likely corresponds to the consolidation of protofibrils into a more ordered helical architecture, representing a transition toward a lower entropic state (79). In the four mutants examined here, however, this maturation pathway is disrupted. Rather than entering productive radial compaction, their ULFs and short filaments undergo extensive aggregation into bundled filaments or non-IF structures. This behavior suggests that the initiation of radial compaction is not merely a passive consequence of filament growth, but a critical mechanistic control step that determines whether assembly proceeds toward mature filaments or is diverted into the formation of aberrant aggregates.

This interpretation is further supported by experiments with fluorescently labeled vimentin, for which ULF formation becomes progressively delayed as the proportion of labeled monomers increases (80). Even modest perturbations of filament subunit structure can therefore shift the balance between productive assembly and defective maturation. In the case of the desmin mutants examined here, early assembly defects appear to lead to a second-stage failure of filament stabilization. This ultimately redirects the system toward non-canonical supramolecular structures.

The transition from filament assembly to aggregation cannot be adequately described within the Rayleigh-Gans framework of static light scattering. Thus, we developed two numerical simulation strategies to capture the two principal classes of aggregate structures formed by the mutants. For the R406W-desmin mutant, which exhibits a phenotype dominated by filament-based aggregation, we employed a worm-like chain model. For the other three mutants, which form more irregular globular complexes, we applied fractal cluster simulations. These models enable us to convert scattering signals into two-dimensional structural representations that can be directly compared with AFM images.

When we parameterized the simulations using persistence lengths and fractal dimensions extracted from the scattering data, they reproduced the experimentally observed morphologies with striking fidelity. This agreement supports the view that the two modeling approaches capture the dominant physical principles underlying the two major modes of mutant aggregation. Importantly, the simulations also highlight the strong mutant specificity of aggregate formation. The mutants differ not only in the extent of aggregation but also in the physical pathway by which aggregation emerges from defective assembly. This distinction is likely to be highly relevant for understanding the heterogeneity of desmin-related disease mechanisms.

Assembly data obtained from mixtures of wild-type and mutant tetramers further support this conclusion. Wild-type desmin can partially offset the effects of mutations such as R406W and L345P, by slowing the kinetics and reducing the extent of aggregate formation. However, it does not abolish aggregate formation entirely. Even in the presence of wild-type protein, smaller and more slowly forming aggregates persist. This indicates that mutant-containing filaments retain abnormal surface properties. These altered surfaces may represent a ‘gain-of-function’ state that could promote inappropriate intermolecular contacts and thereby contribute directly to pathogenicity in patient muscle.

To place these biophysical findings into a physiological context, it is instructive to consider knock-in mouse models of the desmin mutations R406W and R350P, which correspond to R405W and R349P in mice. These models reproduce important aspects of the human disease phenotype and have provided valuable insight into the consequences of mutant desmin expression *in vivo* (51, 81, 82). Additionally, isolated muscle fibers and stable cell lines derived from these mice are useful systems for analyzing altered mechanical and physiological properties associated with the mutant proteins (83–86). The effects of the mutants are evident through altered myocyte structures, changes in passive steady-state elasticity, the progressive disintegration of extracellular matrix fibers, and accelerated muscle fiber aging.

Beyond altered mechanics, the R349P model has revealed a pronounced disturbance of protein homeostasis, leading to substantial desmin aggregate accumulation in muscle tissue (87, 88). An additional unresolved question in the case of desminopathies is whether a defective desmin filament system perturbs mechanotransduction and downstream gene regulation in muscle cells. This is entirely plausible given desmin’s role in connecting the contractile apparatus with nuclei, mitochondria, Z-discs, intercalated discs, and costameres, especially in cardiac muscle (11). Defects in this integrated cytoskeletal network could therefore have consequences that extend well beyond aggregate formation itself.

In summary, the four mutants examined here exhibit distinct defects in filament assembly, aggregation pathways, and underlying structural perturbations. This mechanistic heterogeneity suggests that a single pharmacological strategy is unlikely to be effective across all desminopathies. Instead, successful therapeutic approaches will probably need to address the biophysical pathways specific to each mutation by which normal filament assembly is diverted toward pathological aggregate formation.

## DATA AND CODE AVAILABILITY

Data and python scripts are available on request.

## AUTHOR CONTRIBUTIONS

H.H., B.F., N.M., L.S., S.V.S., and S.K. conceptualized the study. N.M. developed the DWSF method and built the experimental setup. D.S., H.H., and L.S. generated the recombinant protein. L.S. performed and analyzed the DWSF and AFM experiments. J.-P.B. performed and analyzed the SAXS experiments. M.V.D.H. performed and analyzed the MD simulations. L.S. performed the WLC and dimensionality analyses, including their visualizations. L.S., H.H. and B.F. wrote the manuscript. S.K., M.V.D.H., and S.V.S. reviewed and edited the manuscript.

## DECLARATION OF INTERESTS

The authors declare that they have no conflicts of interest with the contents of this article.

## ACKNOWLEDGMENTS

We thank Prof. Dr. Sabine Maier for providing access to the AFM MFP-3D. The authors gratefully acknowledge technical support by Susanne Bauch, Kamila Sabagh and Christian Kuster. Grants from the German Research Foundation (DFG) to Sarah Köster (KO 3572/7-1) and to Harald Herrmann (HE 1853/15-1) are acknowledged. Sergei V. Strelkov received support from the Research Foundation - Flanders (FWO) grant G035223N. L.S. gratefully acknowledges funding of the Erlangen Graduate School in Advanced Optical Technologies (SAOT) by the Bavarian State Ministry for Science and Art.

## SUPPORTING CITATIONS

References (49, 50, 89) appear in the supporting material.

## SUPPORTING MATERIAL

**Table S1:** Results of SAXS measurements. *q*-ranges of the Guinier fit to scatter intensities, mean cross-sectional radii of gyration *RC* (mean ± SD), and apparent filament diameter *DSAXS* for human and mouse desmin assembled at the respective salt concentration and measured by SAXS. *n* denotes the number of independent measurements. These data are displayed in Figure 2 B. Representative SAXS intensity profiles for both species are displayed in Figure S5.

|  | Salt conc. (mM) | 0 | 20 | 40 | 60 | 80 | 100 |
| --- | --- | --- | --- | --- | --- | --- | --- |
| human desmin | q-fit range [$\text{nm}^{-1}$] | [0.132-0.279] | [0.092-0.230] | [0.092-0.201] | [0.052-0.189] | [0.080-0.189] | [0.069-0.189] |
| | $R_C$ [nm] | $3.49 \pm 0.2$ | $4.32 \pm 0.35$ | $4.85 \pm 0.37$ | $5.25 \pm 0.01$ | $5.39 \pm 0.6$ | $5.15 \pm 0.03$ |
| | $D_{\text{SAXS}}$ [nm] | $9.9 \pm 0.6$ | $12.2 \pm 1.0$ | $13.7 \pm 1.0$ | $14.9 \pm 0.03$ | $15 \pm 2$ | $14.6 \pm 0.1$ |
|  | n | 5 | 5 | 9 | 2 | 4 | 2 |
| mouse desmin | q-fit range [$\text{nm}^{-1}$] | [0.089-0.345] | [0.086-0.218] | [0.080-0.195] | [0.075-0.190] | [0.103-0.189] | [0.080-0.178] |
| | $R_C$ [nm] | $2.90 \pm 0.13$ | $4.52 \pm 0.12$ | $5.03 \pm 0.08$ | $5.15 \pm 0.03$ | $5.28 \pm 0.10$ | $5.58 \pm 0.12$ |
| | $D_{\text{SAXS}}$ [nm] | $8.2 \pm 0.4$ | $12.8 \pm 0.4$ | $14.2 \pm 0.3$ | $14.6 \pm 0.1$ | $14.9 \pm 0.3$ | $15.8 \pm 0.4$ |
|  | n | 3 | 3 | 6 | 3 | 3 | 6 |

**Table S2:**
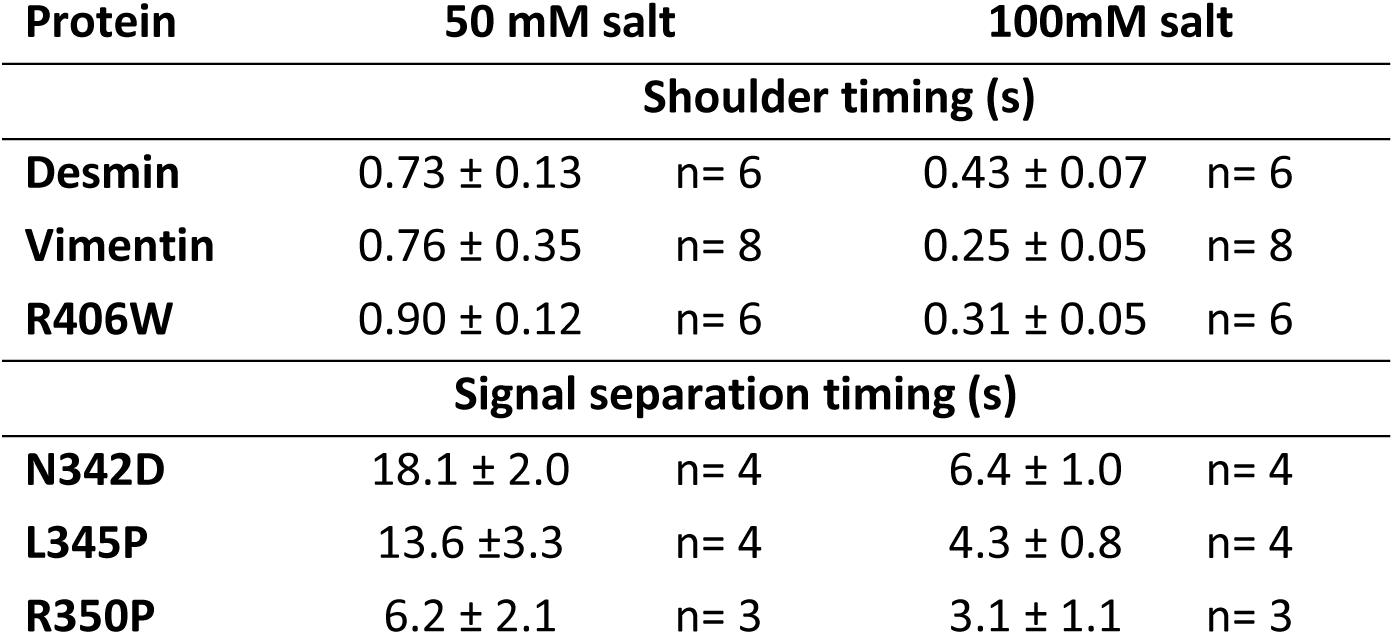
Time points of the characteristic shoulder (mean ± SEM) in the DWSF signal intensities for desmin, vimentin, and R406W-desmin assembly, and time points of the separation (mean ± SEM) of the 405 nm and 594 nm intensity signals for the desmin mutants N342D, L345P, and R350P.

**Figure S1:**
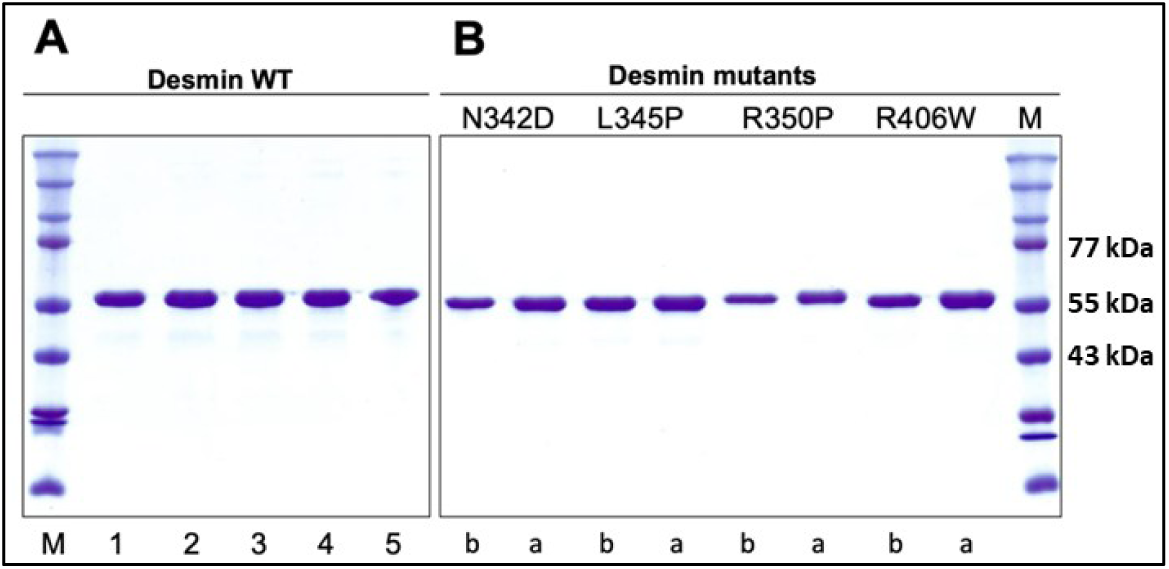
Protein stability documented by gel electrophoretic analysis of recombinant proteins. (A) Wild-type desmin was analyzed. Lane 1: before dialysis; lane 2: after 16h of dialysis into tetramer buffer; lane 3-5: further incubation and at room temperature for 1, 2 and 3 d. (B) Corresponding analysis for the four mutant desmins indicated above the gel: b, before and a, after dialysis. M: Synthetic molecular weight markers (in kilodalton, kDa). Relevant molecular weights are indicated at the right.

**Figure S2:**
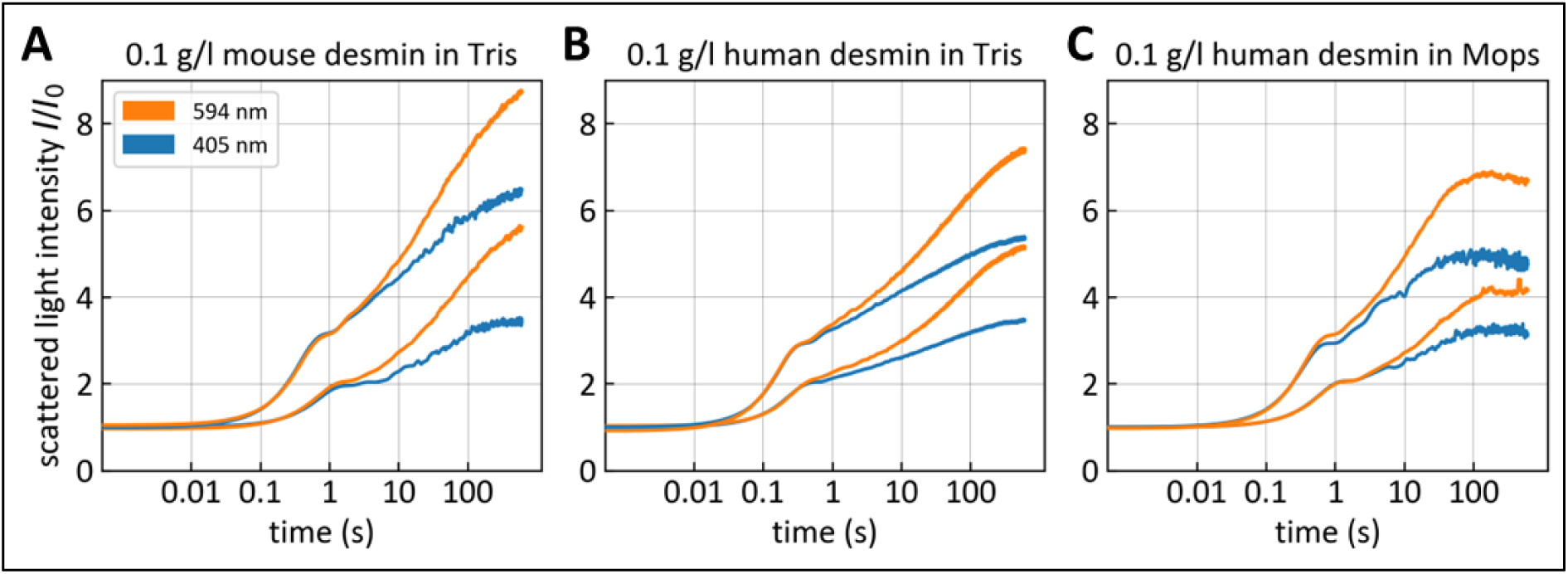
DWSF light scattering profiles of 0.1 mg/ml mouse (A) and human (B, C) desmin in either (A, B) 22.5 mM Tris-HCl buffer (pH 7.5) or (C) 2 mM MOPS buffer (pH 7.5).

**Figure S3:**
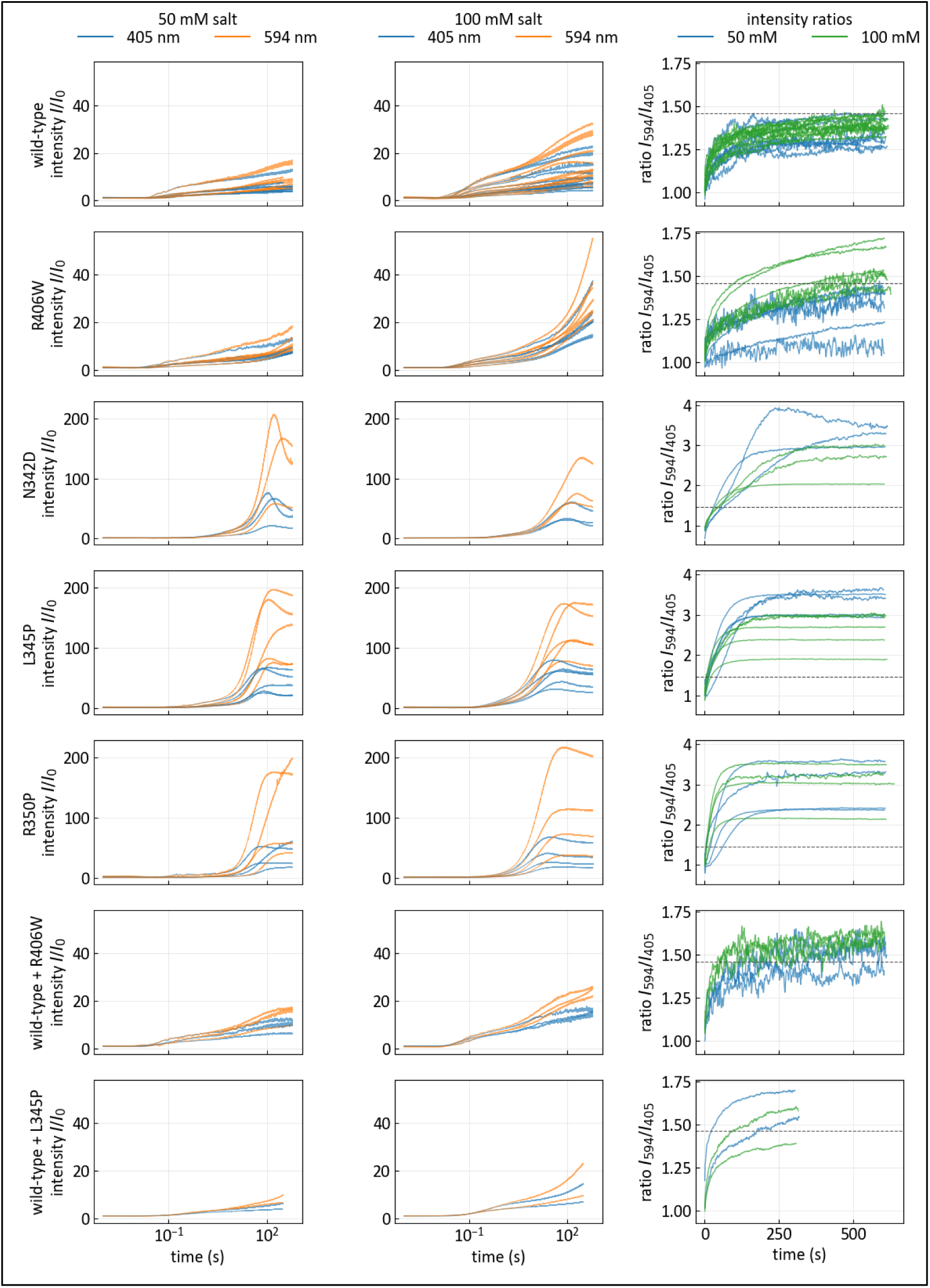
Overview of all measured DWSF light scatter intensities of 0.1 mg/ml desmin as wild-type, mutated variant or wild-type mutant mixture. Assembly in 50 mM (left column) and 100 mM (middle column) salt. Scatter intensity ratios (right column) for the 50 mM salt (blue) and 100 mM salt condition (green). Desmin wild-type and L345P mixed measurements are taken over 5 min assembly time, other proteins over a period of 10 min.

**Figure S4:**
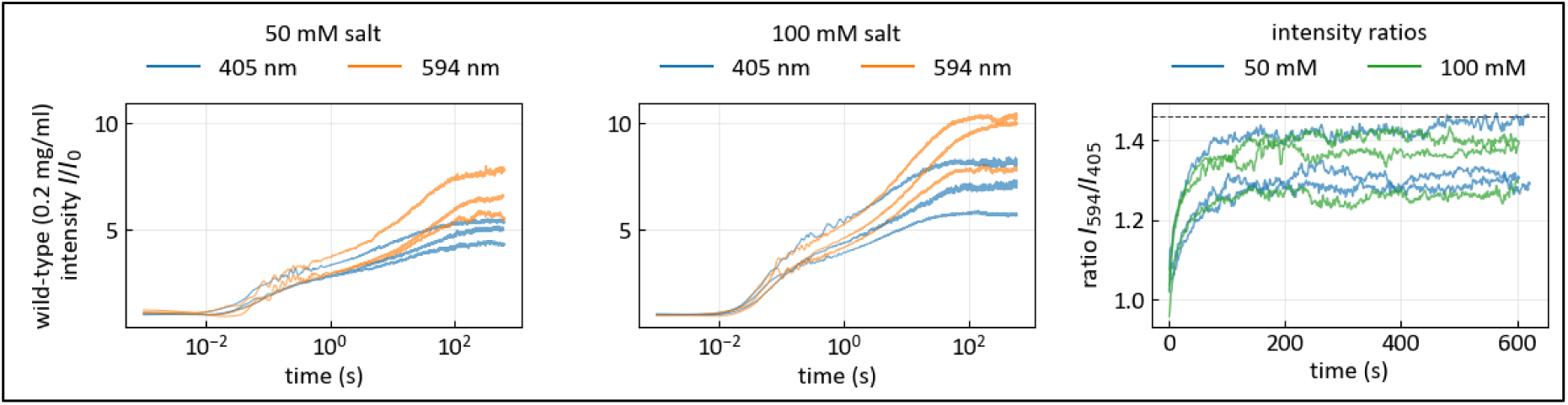
DWSF light scattering intensities for 0.2 mg/ml desmin wild-type assembled at 50 mM (left) and 100 mM salt (middle). Corresponding scatter intensity ratios (right) for the 50 mM salt (blue) and 100 mM salt condition (green).

**Figure S5:**
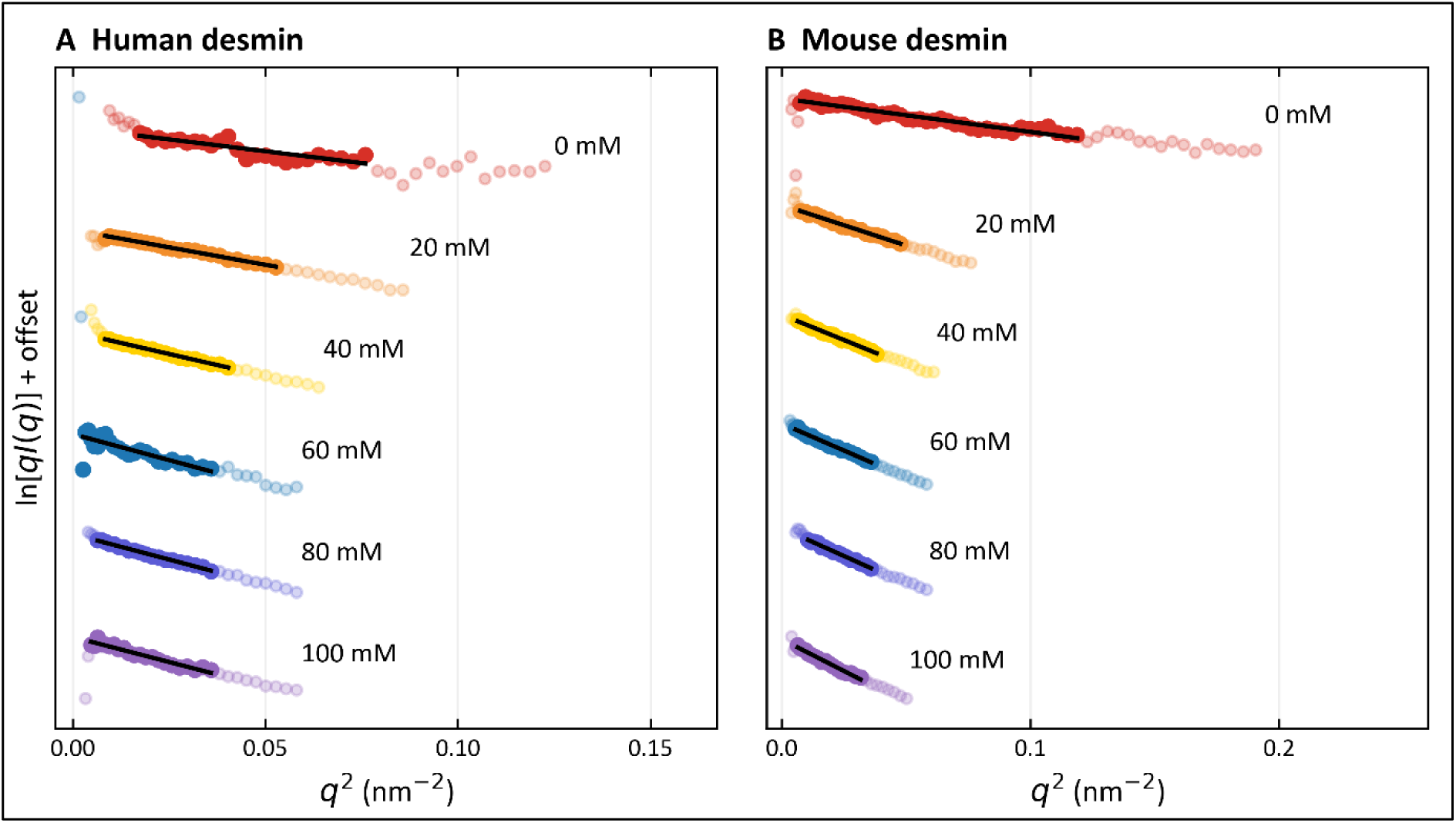
Cross-sectional Guinier plots of representative SAXS profiles for human **(A)** and mouse **(B)** desmin (0.34 mg/ml), assembled at the indicated KCl concentrations. The profiles are shown as *ln[q I(q)]* versus *q^2^* and vertically offset for clarity. Filled symbols indicate data included in the fit, pale symbols show neighboring data outside the fitted interval, and black lines represent weighted linear fits. The cross-sectional radius of gyration *RC* is obtained from the slope according to *m = -RC^2^ /2.* Fitting ranges and corresponding mean RC values of all individual measurements are given in Table S1.

**Figure S6:**
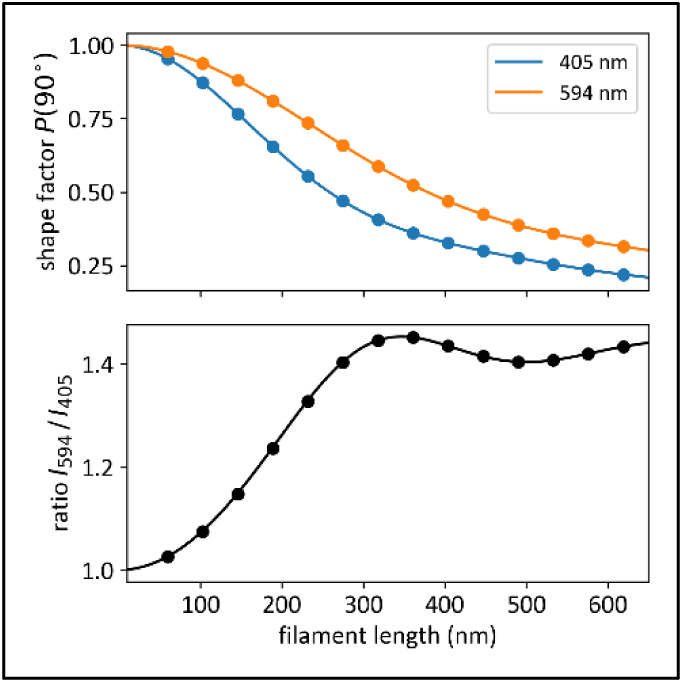
(Upper panel) Shape factor for different wavelengths for scattering angle of 90° versus filament length in nm as predicted by Rayleigh-Gans theory (49). The shape factor determines the normalized scattered light intensities for different filament lengths. (Lower panel) The scattered light intensity ratio I594/I405 increases monotonically during filament elongation up to a filament length of 330 nm (∼7 ULFs). Adapted from (50).

**Figure S7:**
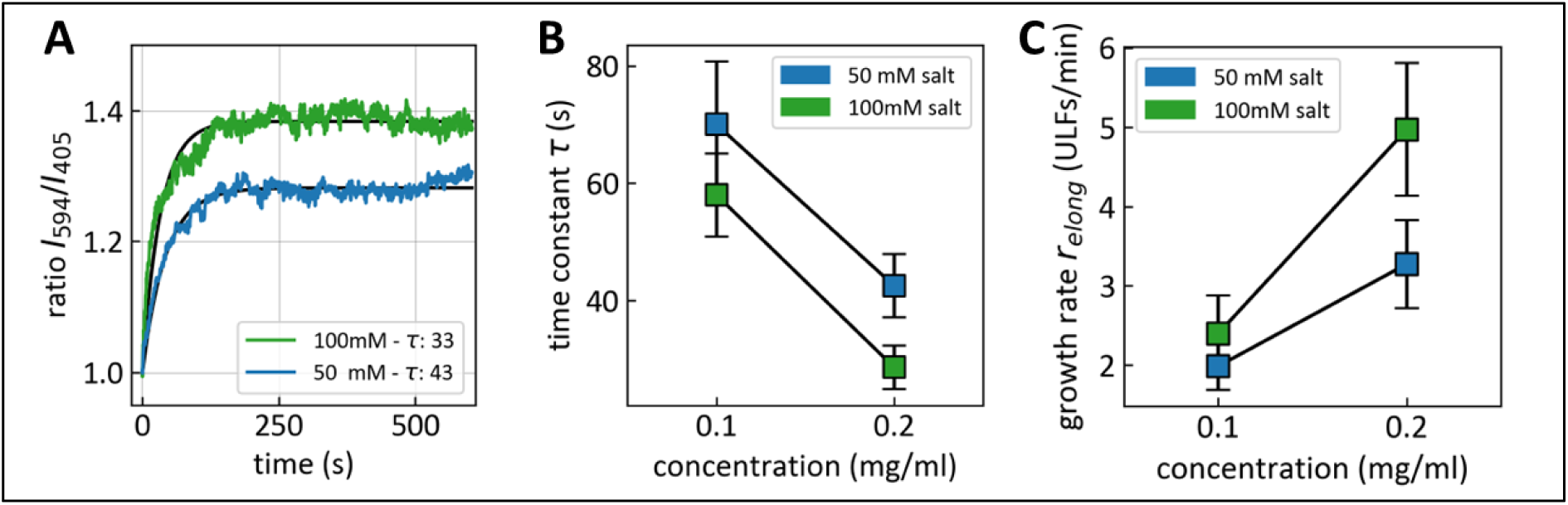
**(A)** Typical scattered light intensity ratio I_594_/I_405_ of 0.2 mg/ml wild-type desmin at 50 mM KCl (blue) and 100 mM (green). Black line indicates an exponential fit *f(t) = 1 + A * (1 - exp(-t/τ))* to the ratio curves, with time constant τ in seconds displayed in the legend. **(B)** Median time constants τ of exponential fits (+-SEM) to scattered light intensity ratios I_594_/I_405_ of 0.1 mg/ml and 0.2 mg/ml at 50 mM or 100 mM KCl in 2 mM Mops buffer or NaCl in 22.5 mM Tris-HCl buffer. **(C)** Filament growth rate of the longitudinal assembly in ULFs/min. Values are calculated from time constants τ shown in (B) and converted to growth rates using the relation *relong = 2.32 / τ*, as reported in (50).

**Figure S8:**
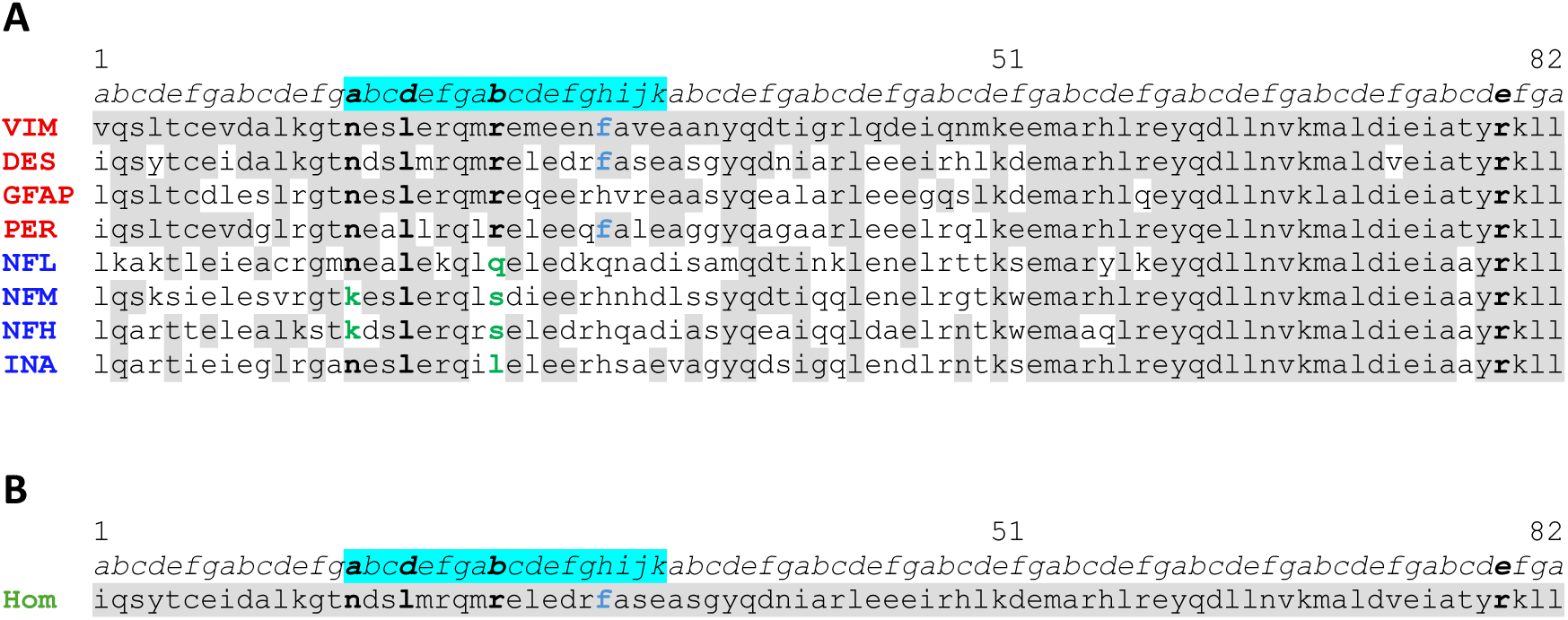
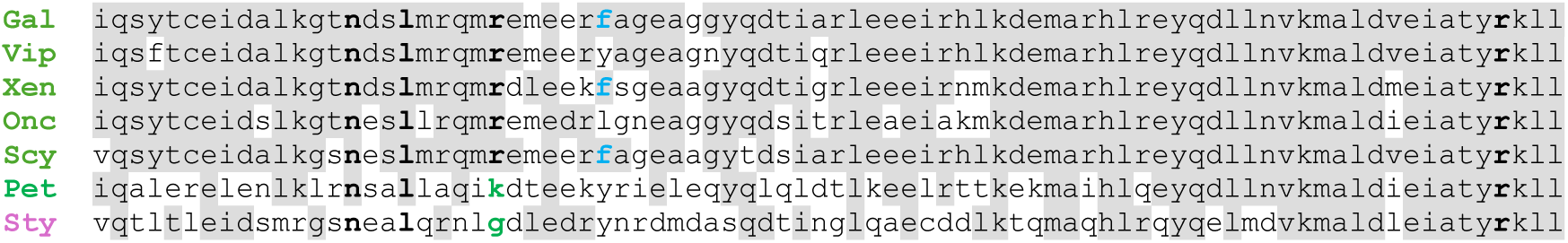
**(A)** Conservation of the desmin amino acids asparagine (342), leucine (345), arginine (350), and arginine (406) in the coil 2 sequences of the human IF proteins constituting sequence homology class III and IV. Sequence comparison of the second half of coil 2 of type 3 (red) and type 4 (blue) human IF proteins to vimentin: VIM, vimentin; DES, desmin; GFAP, glial fibrillary acidic protein; PER, peripherin, NFL, neurofilament light polypeptide; NFM, neurofilament medium polypeptide ; NFH, neurofilament heavy polypeptide ; INA, alpha-internexin. Above the amino acid sequences, the heptad positions are indicated (*abcdefg*), the hendecad repeat is marked in cyan. Amino acids identical to vimentin are shaded gray. The amino acids referring to the mutated desmin aspartic acid (342 - asparagine), proline (345 - leucine), proline (350 - arginine), and tryptohan (406 - arginine) are in bold, black text. Different amino acids are in bold green. The phenylalanine f, which historically marks the “stutter”, is in bold blue. The sequences used are: VIM, GenBank: AAH66956.1; DES, GenBank: AAH32116.1; GFAP, UniProtKB/Swiss-Prot: P14136.1; PER, NCBI NP_006253.2; NFL, UniProtKB/Swiss-Prot: P07196.3; NFM, UniProtKB/Swiss-Prot: P07197.3; NFH, UniProtKB/Swiss-Prot: P12036.5; INA, GenBank: KAI4077333.1. (**B**) Corresponding comparison of the conservation of the desmin amino acids asparagine (342), leucine (345), arginine (350), and arginine (406) in the coil 2 sequences of typical vertebrates (green) and one invertebrate (pink). Sequence comparison of the second half of coil 2 of desmin from human (Hom), chicken (Gal), *vipera berus* (Vip), *Xenopus laevis* (Xen), *Oncorhynchus mykiss* (onc), *Scyliorhinus stellaris* (Scy), *Petromyzon marinus* (Pet), *Styela plicata* (Sty). The sequences used are in the order of their use: Homo sapiens, GenBank: AAH66956.1.; Gallus gallus (chicken), NCBI NP_001383608.1; Vipera berus (crossed viper), NCBI XP_081169529.1; Xenopus laevis (clawed toad), NCBI NP_001080177.1; Oncorhynchus mykiss (rainbow trout), NCBI XP_036831228.1; Scyliorhinus stellaris (nursehound, shark), GenBank: CAC83054.1; Petromyzon marinus (sea lampray), NCBI XP_032828666.1; Styela plicata (pleated sea squirt); GenBank: CAA06293.

**Figure S9:**
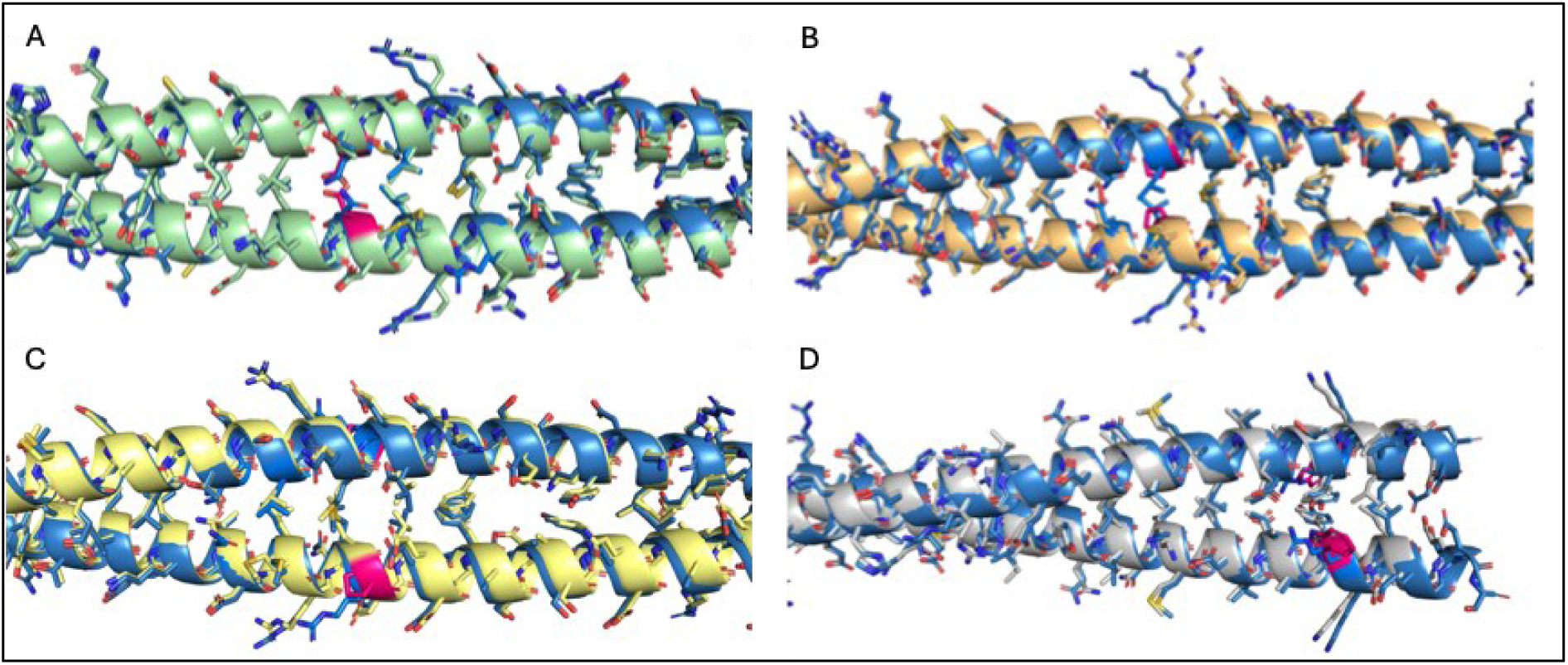
Superposition of desmin coil 2 predicted by AlphaFold3 with its mutant variants (89). The wild-type structure is shown in blue; panels A–D depict mutants N342D, L345P, R350P, and R406W, respectively. The mutated residues are highlighted in pink.

## Notes

### Competing Interest Statement

The authors have declared no competing interest.

### Summary of Updates

Updated Method and Result section on SAXS data. Additional Supplementary Figures S2-S5 showing complete DWSF scatter data and background information.

